# Local adaptation of the mosquito vector, *Aedes aegypti,* and implications for predicting the effects of temperature and climate change on dengue transmission

**DOI:** 10.1101/2025.02.14.637444

**Authors:** Nina L. Dennington, Marissa K. Grossman, Janet L. Teeple, Leah R. Johnson, Marta S. Shocket, Elizabeth A. McGraw, Matthew B. Thomas

**Affiliations:** Department of Entomology, The Pennsylvania State University, University Park, PA, USA; The Center for Infectious Disease Dynamics, The Huck Life Sciences, The Pennsylvania State University, University Park, PA, USA; Department of Statistics, Virginia Tech, Blacksburg, VA, USA; Department of Geography, University of Florida, Gainesville, FL, USA; Lancaster Environment Centre, Lancaster University, Lancaster, UK; Department of Biology, The Pennsylvania State University, University Park, PA, USA; Department of Entomology & Nematology, University of Florida, Gainesville, FL, USA; Invasion Science Research Institute, University of Florida, Gainesville, FL, USA; Department of Biology, University of York, York, UK

**Keywords:** Mosquito, adaptation, Temperature, Climate change, Ecology, Population fitness, Vector-borne disease transmission, Thermal performance

## Abstract

There is concern that increases in temperature due to climate change could lead to shifts in the dynamics and distribution of mosquito vectors. Many current models assume there are ’average’ thermal performance curves for a given vector species transmission. However, this ‘one-size-fits-all’ assumption ignores the potential for local adaptation to create population-specific differences in thermal performance. In this study, we explored thermal performance of five independent field populations of *Ae. aegypti* from Mexico, together with a standard laboratory strain. We reared these six populations at temperatures between 13°C-37°C to generate thermal performance curves for a suite of life-history traits. Composite models integrating these traits revealed the effects of temperature on population growth rates and dengue virus transmission potential. The results provide strong evidence for the potential for local adaptation in *Ae. aegypti* populations, challenging applicability of ‘one-size-fits-all’ thermal performance models to assess climate impact on mosquito-borne diseases.

## Introduction

Dengue is the most widespread vector-borne disease of humans, with more than 3.9 billion people estimated to be at risk in over 128 countries (1). Dengue virus is transmitted by mosquitoes, with *Aedes aegypti* as the principal vector (2). Transmission of dengue virus is strongly climate-sensitive (2–5) and as such, there is substantial interest in the extent to which disease dynamics will be affected by climate change (6). Numerous studies suggest a likely net increase in the overall size of the population at risk of dengue and other arboviruses in the coming decades (7–14).

The climate-sensitivity of mosquito-borne diseases derives from the fact that many mosquito and pathogen life history traits are strongly affected by temperature. Traits of many ectotherms are temperature-dependent, often exhibiting nonlinear thermal performance curves (15–17). For mosquitoes and their pathogens, these traits include both physiological traits, such as larval survival, development rate, adult survival, fecundity, vector competence, and pathogen development rate, as well as behavioral traits such as biting rate, mating and foraging behaviors (10,11,18–30). Additionally, these traits show variation between individuals depending on nutritional status, body size, and genetics (31–34).

Thermal performance curves provide a framework to assess how environmental temperature affects biochemical and physiological processes. In order to understand how changes in vital rates impact transmission, it is necessary to integrate individual traits into temperature-dependent fitness metrics such as the Basic Reproduction Rate (R_0_, defined as the number of secondary cases resulting from an initial primary case introduced into a population of susceptible hosts), or vectorial capacity (a measure of the transmission potential of a mosquito population defined as the number of potentially infectious bites that would eventually arise from all the mosquitoes biting an infectious host on a single day). These temperature-dependent transmission metrics have been used extensively to investigate variation in the relative environmental suitability for transmission over time and space (7–11,22,28,35–42).

Common to nearly all mechanistic models that follow the thermal performance-R_0_ approach, is the assumption that thermal performance curves are fixed for a given mosquito species and associated pathogen. However, this assumption ignores the potential for the environment to impact thermal performance curves at the local population level (43). Local adaptation can occur when there is spatial variation in selection due to interactions with the environment leading to a relative increase in fitness at a local level (43). The evidence for local thermal adaptation in insects is mixed. Some studies indicate the potential for populations to shift thermal performance curves through evolutionary or plastic responses (44–49), while others suggest limited adaptation in response to variation in climate (50–52). In general, it is expected that short generation times and high intrinsic population growth rates, both of which are true for mosquito vectors, should increase the probability of adaptation to changing conditions (53,54). Local adaptation in mosquitoes to other environmental drivers, such as insecticide exposure, humidity, or daylength, support this suggestion (54–60).

Common garden experiments, in which populations collected from distinct geographic locations are examined together under shared conditions, have been used to compare thermal performance of different insect populations collected across environmental gradients (32,61–65). Recently, we followed this approach to demonstrate differences in thermal tolerance (measured as knockdown rate in response to exposure to stressful high temperature) between five field-derived populations of *Ae. aegypti* from different locations in Mexico, together with a longstanding lab colony (48). These results were strongly suggestive of local adaptation, but knockdown rate does not describe overall fitness and this single measure does not enable us to explore possible implications for transmission. The aim of the current study, therefore, is to extend research in this system to characterize thermal performance of a suite of life history traits and determine whether there are between-population differences in thermal dependence of mosquito fitness and dengue transmission potential. We find strong evidence for local adaptation, challenging the utility of ‘one-size-fits-all’ thermal performance models to capture the spatial or temporal influence of temperature and temperature change on transmission risk.

## Materials and Methods

Methods used in the current study closely follow those outlined in Dennington et al. 2024 (48).

### Mosquito Collection

*Aedes aegypti* mosquitoes were collected from the field in five different locations in Mexico (Cabo San Lucas, Acapulco, Monterrey, Ciudad Juárez, and Jojutla) using ovitraps. Populations from Mexico were founded with at least 100 females and to remove the influence of maternal effects, mosquitoes were reared in the lab for at least one generation in standard laboratory conditions (27°C, 80% humidity, 12:12hr photoperiod) prior to experimentation. Two populations, Jojutla and Ciudad Juárez, were reared for an additional generation to ensure a large enough population for subsequent experiments. The field locations were chosen to capture a gradient of the climate and landscape (see S1 Table). The field populations were compared to a standard laboratory population (Rockefeller strain) that were maintained at Penn State University under standard insectary conditions over many years.

### Experiments to Generate Temperature-Dependent Data

Mosquito life-history traits including egg-to-adult survival, mosquito development rate, mean adult survival, fecundity, and biting rate were measured in mosquitoes reared at 13°C, 15°C, 19°C, 23°C, 25°C, 27°C, 29°C, 31°C, 33°C, 35°C, 37°C, each ± 0.2°C and 80 ± 10% relative humidity in environmentally controlled incubators. These life-history measurements were replicated three times at each temperature for each population. We began with eggs from the five field populations and one laboratory line were hatched at 27°C for 24 hours. Then, 200 first instar larvae were put into 1.89 L containers with 1 L of deionized water and 0.20 mg of larvae bovine liver powder (MP Biomedicals) and each of the three replicates were placed in the incubator at their respective temperatures. We fed larvae 0.20 mg of liver powder per larvae every other day until pupation, but once pupation began we scaled their food to the remaining number of larvae. Pupae, both living and dead, were removed and counted on the day of pupation and placed in a small cup (30 mL) with water from their original environment to allow for eclosion. Pupae were then added to a small cage (17.5 cm^3^) with continuous access to 10% sugar solution (dextrose anhydrous and deionized water). We counted the number of adults that eclosed every day. After 95% of surviving females emerged, we blood-fed females after 3-5 days. We used blood from de-identified human donors (BioIVT, Corp.) and so IRB approval and human subjects’ approval was not needed. We immediately counted the total number of blood-fed females and placed up to 10 individual females into separate containers (50 mL polypropylene centrifuge tubes) that were lined with filter paper and 7 mL deionized water to measure individual fecundity. We also placed up to 20 females into two small cages (10 in each) with a small filter paper for egg laying to monitor adult survival. We recorded the day that females in individual containers first laid eggs, which we used for fecundity measures and to approximate the biting rate (1/gonotrophic cycle length), after which we removed them from their containers and placed them into the group cages. We extracted the water from the containers to let the filter paper dry in their respective incubators and then we counted the number of eggs from individual mosquitoes. For the course of the experiment, we offered each cage of females a blood meal every 4 days, and counted the number of adults that died every day. We censored this experiment 4 weeks after the first egg lay at each temperature.

### Models of Thermal Performance Curves

To analyze the thermal performance curve data we used a Bayesian approach following methods described in Johnson et al. (2015). We fit a symmetric thermal response function to our data for fecundity (number of eggs per female for the first gonotrophic cycle) and egg-to-adult survival using a quadratic equation:

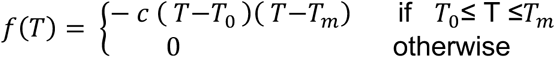

Mosquito development rate and biting rate were described by a Brière function:

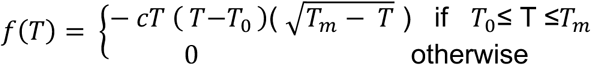

where T_0_ is the thermal minimum (below which the function is zero), T_m_ is the thermal maximum, and c is a positive constant that controls the curvature of the function (and thus the height of the curve for a given value of T_0_ and T_m_). Blood feeding rate was approximated as the reciprocal of the duration of the first gonotrophic cycle, a standard assumption used in many studies (10,66–68).

We assume that observed values for fecundity, mosquito development rate, and biting rate must be non-negative, and are thus were modeled with a truncated normal likelihood:

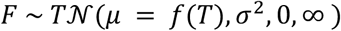

We chose priors for T_0_ and T_m_ to restrict each trait to its biologically realistic range, assuming temperatures below 0°C and above 45°C were fatal as previously measured (10,11,22,37). For other parameters we chose relatively uninformative priors (*SI Appendix*, Supporting Text).

In contrast to fecundity, biting rate, and development rate, egg-to-adult survival is a probability and must be constrained to lie between 0 and 1. Thus, we modeled these data as being binomially distributed:

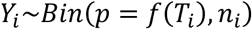

where *n* is the number of total observations of which Y were successes (i.e., survival to become adults) and the probability of a success at a particular temp, *p,* depends on temperature (f(*T*)), specifically being a piecewise quadratic, as above, but being constrained to be less than or equal than 1.

We also measured survival times (life spans) for adult mosquitos by counting the number of mosquitoes that died each day. Time to death for adult mosquitoes in each treatment were observed until a censoring time *T*_cut_; i.e., for each temperature, survival observations were censored 28 days after the first egg lay. Thus, at each temperature for each population most mosquitoes were observed until they died, but a portion of them were still alive at the end of the 28 days, and so the survival times were right-censored. We used a variation of a Bayesian Weibull survival model. We assume that the observed lifetimes, *y*_*i*_, at some particular temperature are drawn from a Weibull distribution with rate parameter β and shape parameter *k*, with pdf,

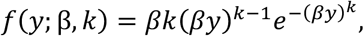

and corresponding CDF notated as *F*(*y*; β, *k*). The median, *m*, of the Weibull is defined as

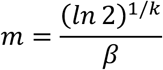

which, can be rearranged to solve for *β*. For interpretability, we assume that the median lifetime decays exponentially with temperature across the experimental temperature range studied, that is,

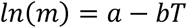

so that the median thermal performance curve, *g*(*T*), is

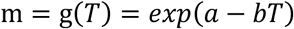

where *T* is the temperature, and *a* and *b* are parameters to be estimated.

The likelihood for the survival data is comprised of two components, one describing the individuals that were observed to die within the study period (δ_i_ = 1), and those who are censored (i.e., that survive the study period, δ_i_ = 0). Thus, the likelihood is given by

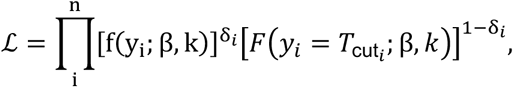

where the rate parameter is defined in terms of the median thermal performance curve, β = 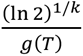.

We fit all models described above using Markov Chain Monte Carlo (MCMC) sampling that is implemented in JAGS, using the R package *R2jags* (69–71). For each life history trait or thermal response, we ran five MCMC chains with a 5,000-step burn-in and saved the subsequent 10,000 steps. We thinned the posterior samples by saving every eighth sample, for a total of 3125 posterior samples of parameters. We used these posterior samples to produce samples from the posterior distribution of each trait across temperature. We then summarize the relationship between temperature and each trait by calculating the mean and 95% highest posterior density interval (HPD interval) for the function across temperature. The HPD interval is a type of credible interval that includes the smallest continuous range containing 95% of the probability, which is implemented in the coda package (72).

### Mosquito Fitness and Transmission Calculations

To characterize overall fitness, we used a temperature-dependent model for population growth *r*(*T_i_*) as previously described in Amaraesekare and Savage (73):

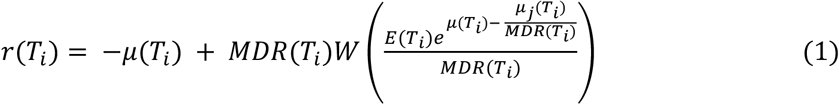

Here MDR is mosquito development rate, E is eggs per female measured at the first gonotrophic cycle, *μ* is adult mortality rate and *μ*_9_ is juvenile mortality rate (*SI Appendix*, S1 Fig, S7 Table). W(x) is the upper branch of the Lambert function. We combined the thermal performance curves for each trait to calculate temperature-dependent fitness, *r*(*T_i_*), creating a unimodal curve. We followed Amarasekare and Savage (73) and truncated the results at r(Ti) = 0 rather than allowing them to go negative.

We estimated the posterior distribution of *r*(*T_i_*) and used it to calculate the key temperature values for relative temperature dependent population fitness. We used the mean and 95% credible intervals (95% CI) for the critical thermal minimum, maximum and optimum temperatures for population fitness.

We used temperature-dependent models of transmission to investigate the potential implications for transmission of dengue virus, utilizing a previously published framework (10,11,22,37). This approach models transmission rate as the basic reproduction rate R_0_, which is defined as the number of secondary infections originating from an initial infection when introduced to a completely susceptible population. The temperature dependent R_0_ is given by:

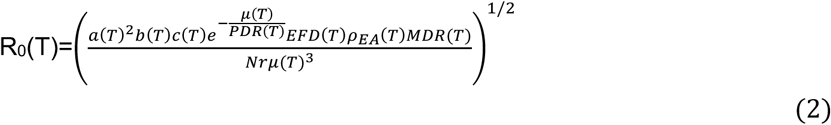

In this equation, (T) indicates that the trait is a function of temperature, a is the biting rate per mosquito, b is the proportion of infectious bites that infect susceptible humans, c is the proportion of bites on infected humans that infect uninfected mosquitoes (*b* × *c*=vector competence), μ is the adult mosquito mortality rate, PDR is the pathogen development rate (the inverse of the extrinsic incubation period), EFD is the number of eggs a female produces each day, *ρ*_*EA*_ is the survival probability of mosquitoes from egg to adult, MDR is the mosquito development rate (the inverse of the egg-to-adult development time), N is the density of humans, and r is the human recovery rate.

In order to use this metric in this study, we have to make two approximations to connect the data and fits obtained from our empirical study to this expression. The first is in the adult mosquito mortality rate. In the original derivation of R_0_, it is assumed that mosquito lifetimes are exponential distributed with rate parameter μ. Thus, under this assumption, the mean lifetime, *λ*, is 1/μ, and the median lifetime, (denoted at *m* above), would be equal to 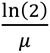. Thus, everywhere in the R_0_ equation we let 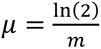. Similarly, we must approximate the mosquito development rate by taking the 1/ mosquito development time to get the rate of development.

In the current study we did not examine infected mosquitoes and so we used previously published thermal performance curves from Mordecai et al. (2017) for vector competence and pathogen development rate for dengue virus in *Ae. aegypti* (10). Also, because we have no measures of human density or recovery rate we follow the approach of Mordecai et al. (2017) and normalize the R_0_ curves to a scale of 0-1. This measure of Relative R_0_ enables comparison of key features of thermal dependency including the critical minimum temperature (CTmin), the critical maximum temperature (CTmax), the optimum temperature (Topt) and the thermal breadth (Tbreadth, defined as the range where performance is above 80% optimal (74,75) but does not provide estimates of maximum performance (Pmax).

#### Testing for Differences Between Populations

We used DIC (Deviance information criterion) to test for statistically significant differences between the thermal performance curves of different populations. Specifically, for each trait, we compared the DIC of a global model fit to the data from all populations to the sum of DIC scores from the models fit separately to data from each population (equivalent to fitting a single model assuming all parameters may vary between populations). Populations were considered significantly different for a given trait if the sum of DIC scores from the separate models was >= 2 DIC units lower than the DIC score from the global model.

## Results

The estimated thermal performance curves for egg-to-adult survival probability, mosquito development rate, fecundity, biting rate, and mean adult survival indicated by both the empirical data and the Bayesian model fits are presented in Fig 1A-E, with the key features of these curves (i.e. the thermal optimal temperature (Topt), the maximal performance (Pmax), the critical thermal minimum (CTmin), critical thermal maximum (CTmax) and thermal breadth (Tbreadth) summarized in the SI (S2-6 Tables).

**Figure 1.**
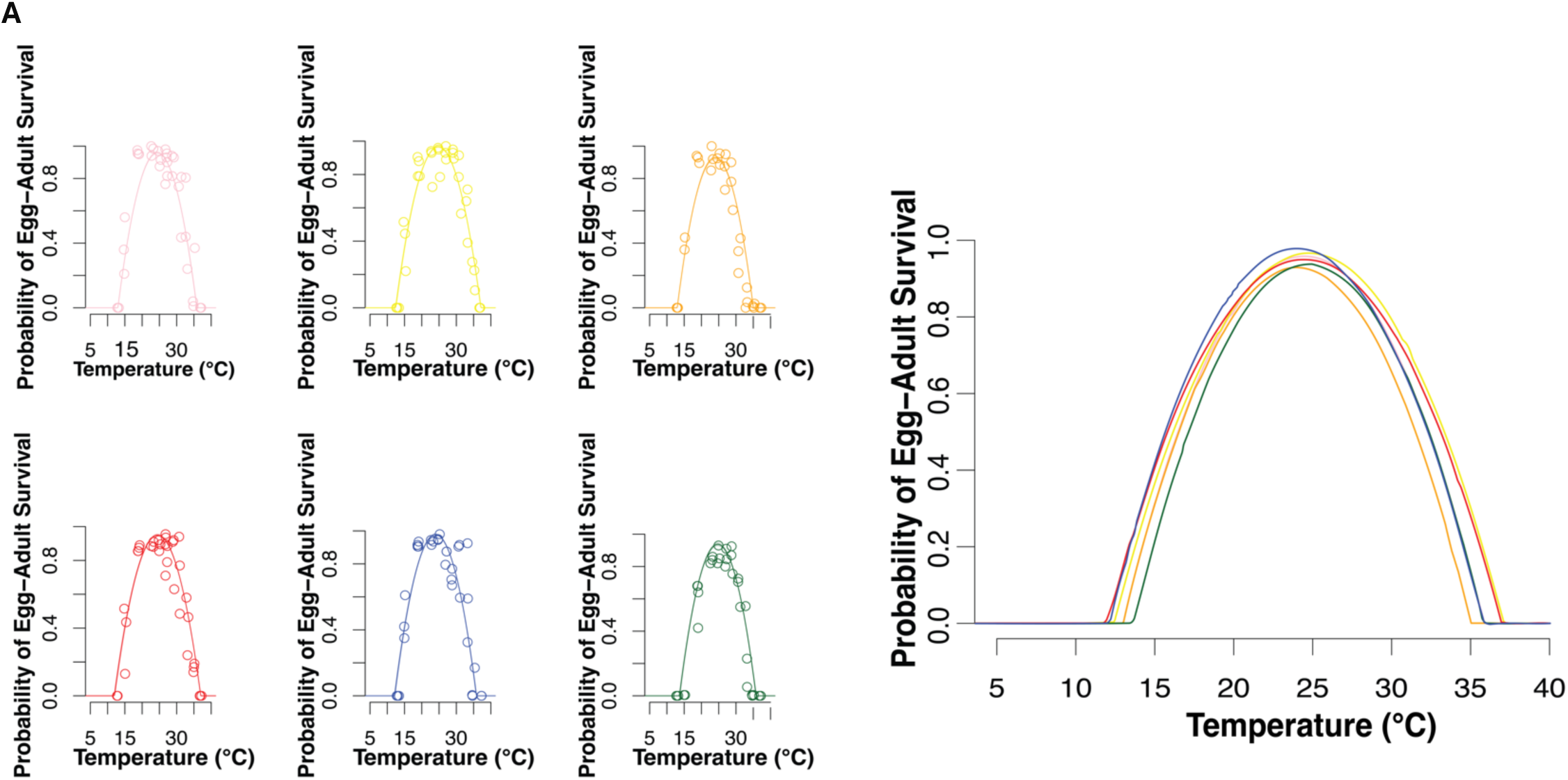
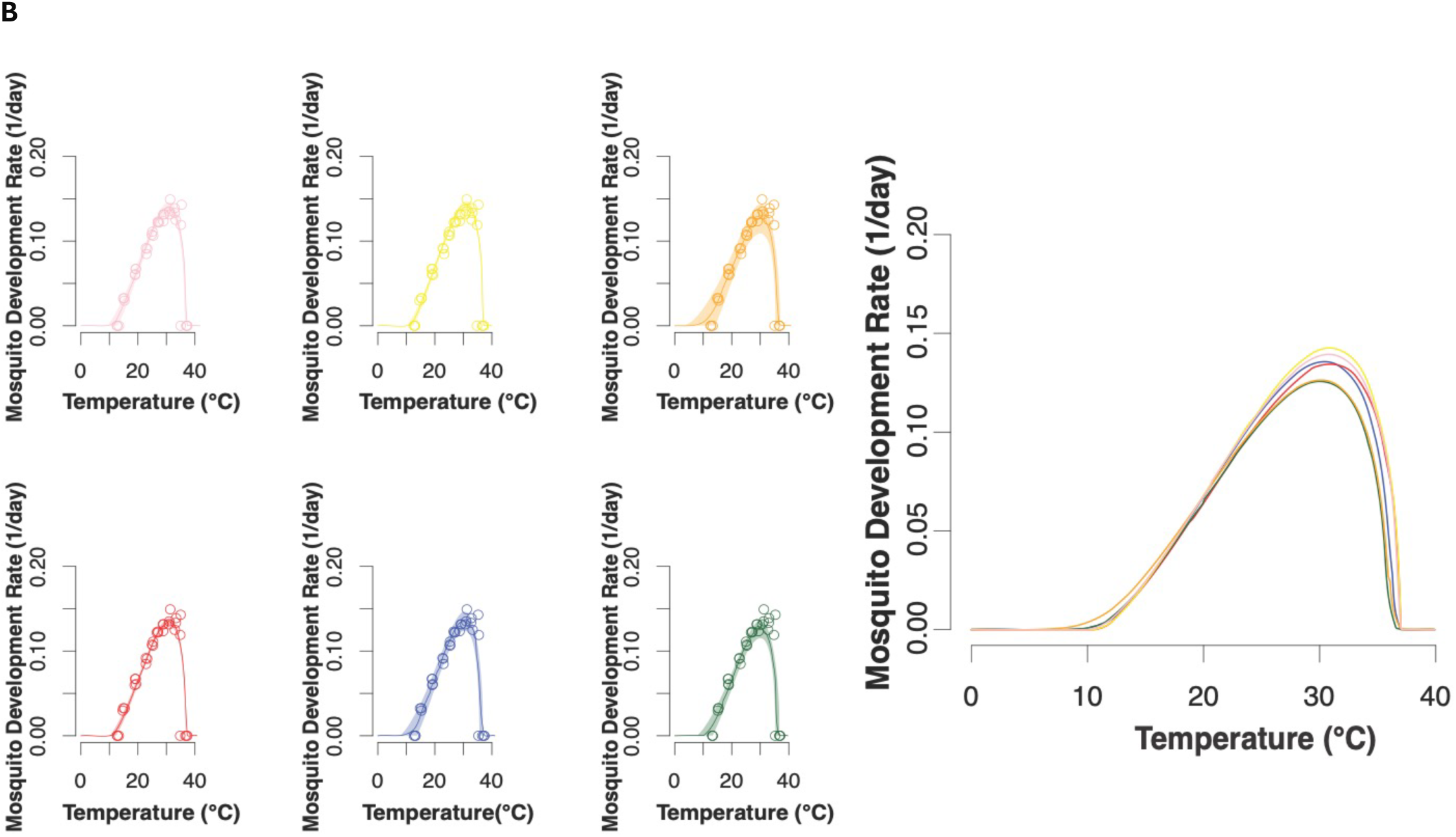
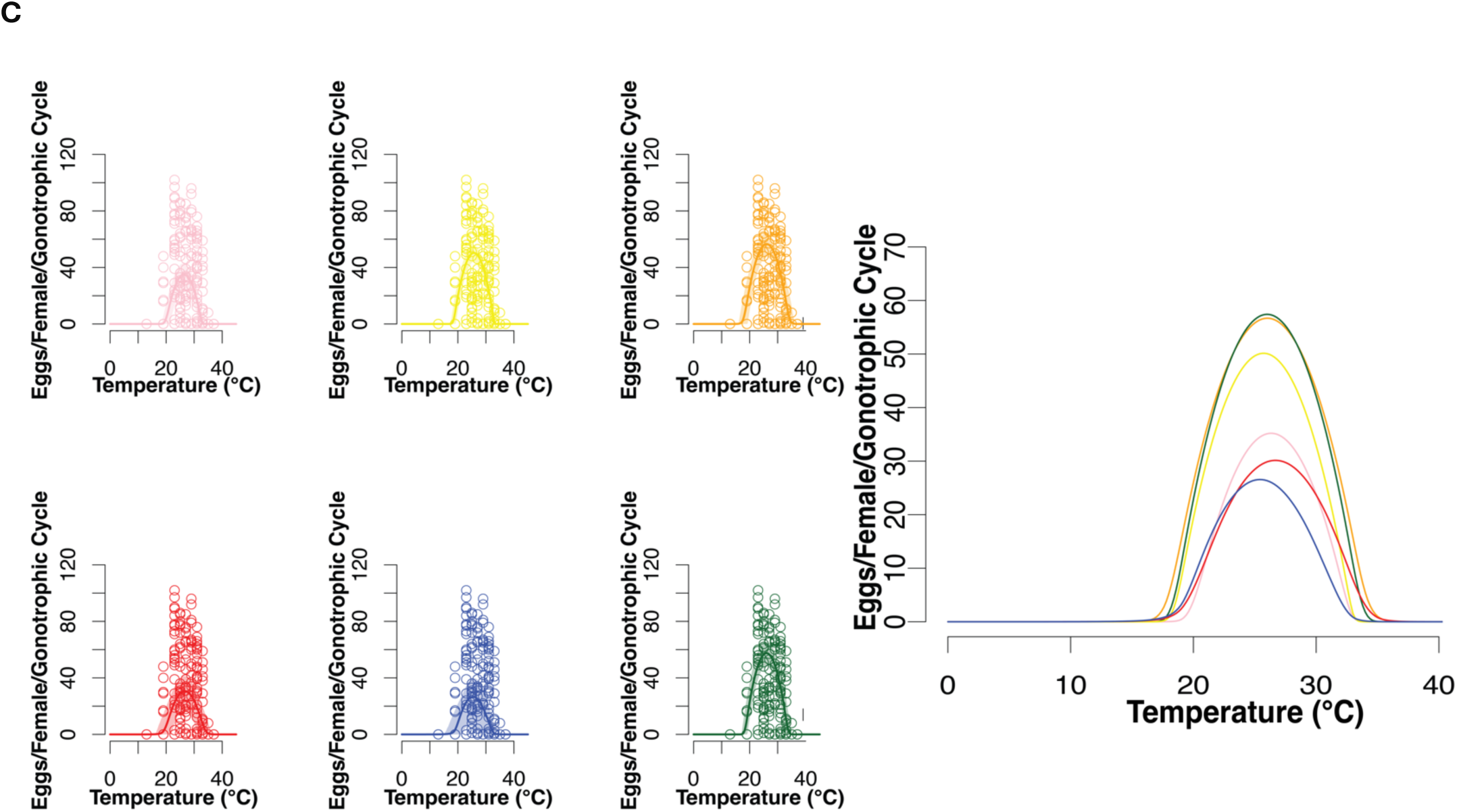
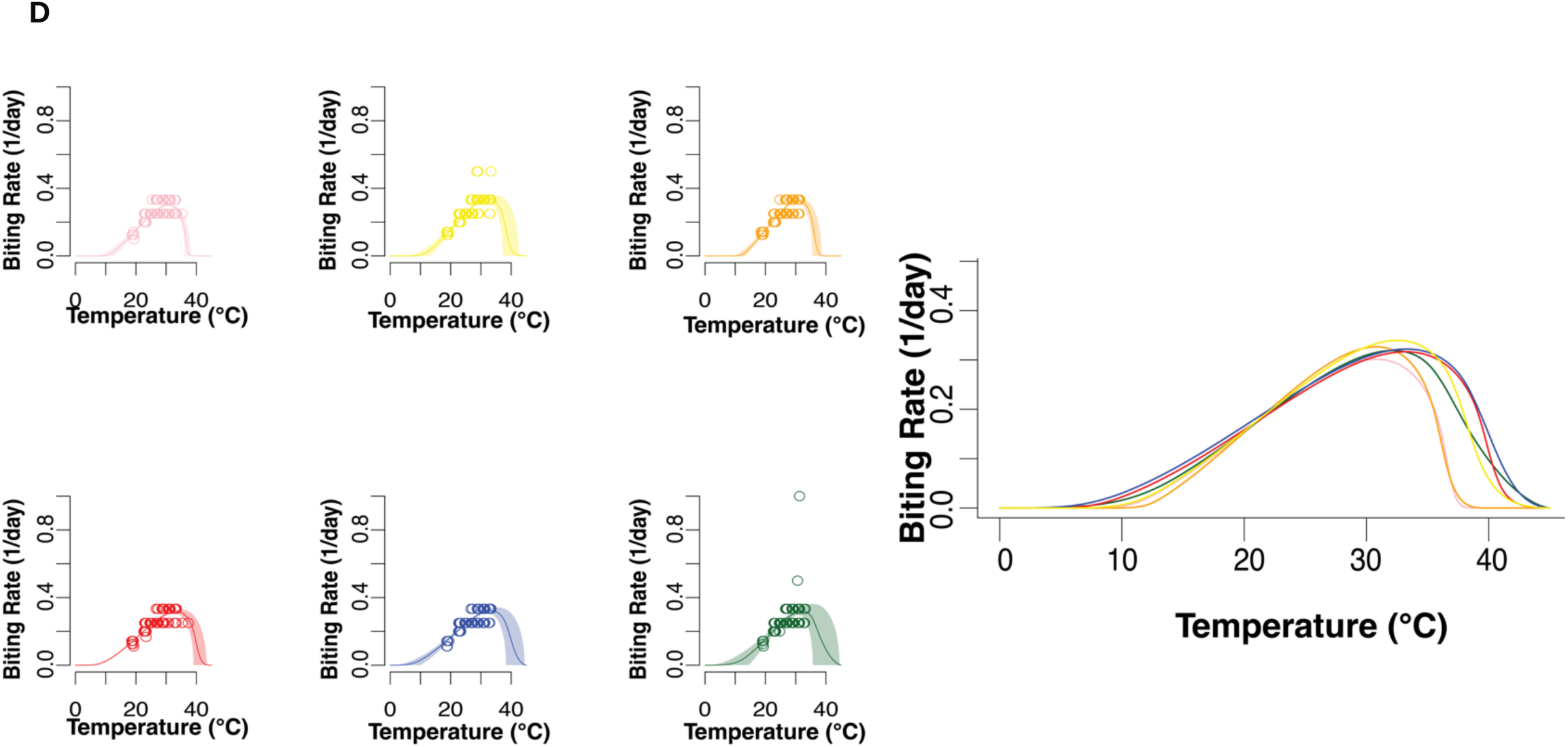
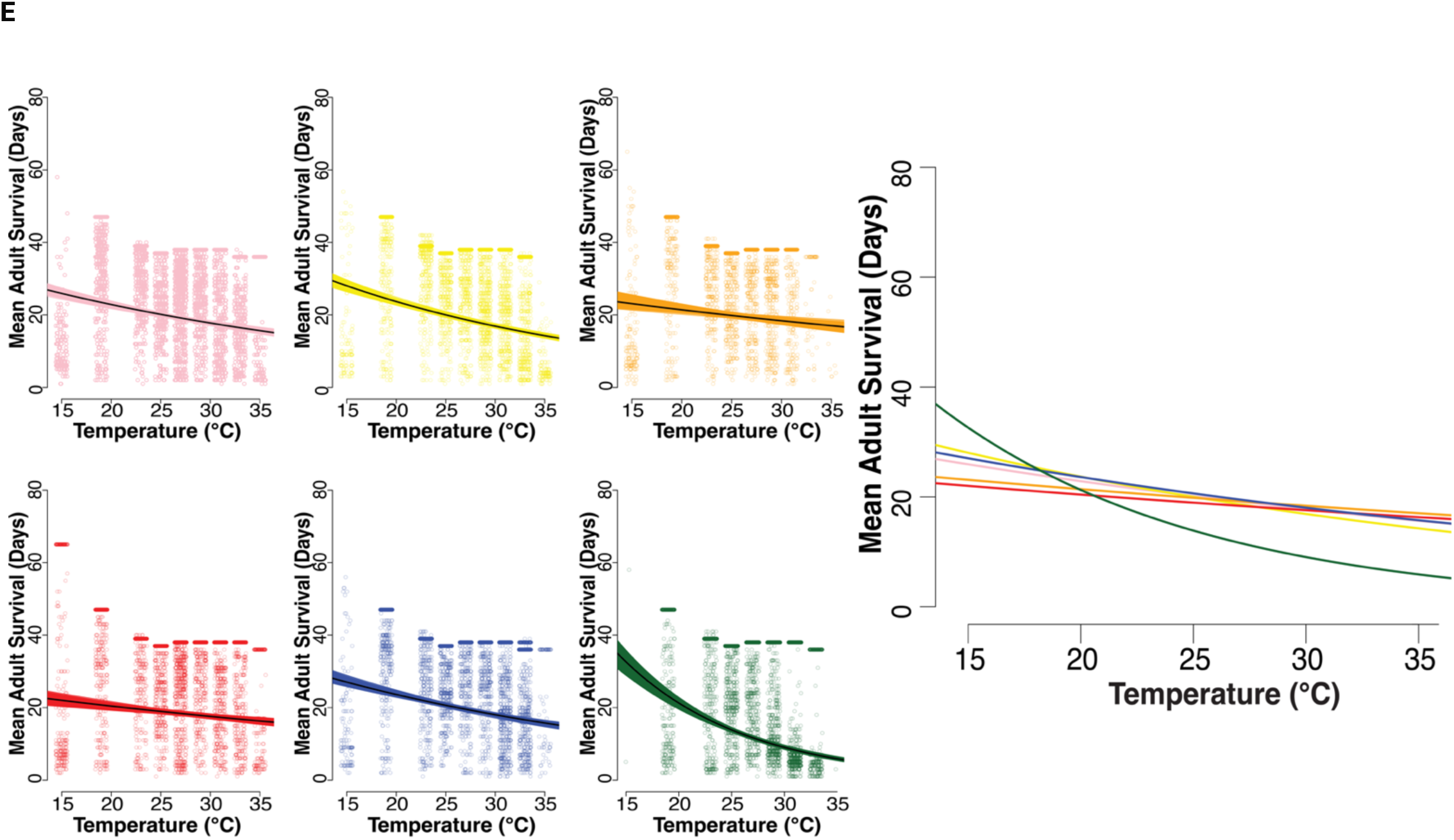
Life history traits measured for *Ae. aegypti* mosquitoes reared at 13°C, 15°C, 19°C, 23°C, 25°C, 27°C, 29°C, 31°C, 33°C, 35°C, 37°C. We measured life history traits for five populations from Mexico (Cabo San Lucas (yellow), Acapulco (pink), Monterrey (orange), Ciudad Juárez (red), and Jojutla (blue)) compared to a laboratory adapted line (shown in green). The data points for figures are the values from three replicates for each corresponding population at each temperature with individual values for adult survival and fecundity along with mean values for each replicate for biting rate, mosquito development rate and egg-to-adult survival probability. Thermal performance curves for life history traits for the six populations were fit using Bayesian inferences with weakly informative priors are shown (Model values can be found in SI Table S2-6). Each mosquito line was tested between 13°C-37°C, but individuals did not survive and reproduce at 13°C and 37°C. **A)** Egg-to-adult survival measured as the probability of individuals surviving to the adult stage. **B)** Mosquito development rate is the inverse of the amount of time that it takes to reach the adult stage. **C)** Fecundity is measured as individual egg production for the first gonotrophic cycle. **D)** Model fits for approximated biting rate (1/gonotrophic cycle length). **E)** Model fits for estimated mean adult survival (in days) derived from measures of daily survival of adults over 28 days post initial blood meal, censored individuals are clustered at the last day tested.

The between-population differences in the thermal performance curves for the juvenile traits of egg-to-adult survival and development rate were relatively small, with 1°C or less for the differences between the respective CTmin, CTmax, Tbreadth values of the different populations, although still significant according to the Deviance Information Criterion (DIC) (Fig 1A, B; S2-S6 Tables). For the adult traits of fecundity, survival and biting rate, the differences were more marked. Specifically, for eggs per female, there was an approximate 2°C difference between the lowest (Monterrey population) and highest (Acapulco population) values for CTmin, and 3°C difference between lowest (Jojutla population) and highest (Monterrey population) values for CTmax (Fig 1C). For the biting rate approximation, there was an approximate 6.6°C difference between the lowest (Jojutla population) and highest (Monterrey population) values for CTmin, and 4.5°C difference between lowest (Monterrey population) and highest (Jojutla population) values for CTmax (Fig 1D). For adult survival, the data did not exhibit a clear unimodal pattern across the temperature range studied so it is not possible to define values for CTmin and CTmax (Fig 1E). Nonetheless, the thermal performance models indicate significant differences between populations according to the DIC criteria. For the field strains, the population from Ciudad Juárez survived the worst at lowest temperatures while the population from Cabo San Lucas survived the best. Alternatively, the population from Cabo San Lucas survived the worst at high temperatures while the population from Monterrey survived the best. Notably, the longstanding lab strain exhibited the most pronounced thermal response, with the highest survival at cool temperatures and the lowest survival at warm temperatures.

The combined effects of variation in individual traits are shown in the thermal performance curves for overall mosquito fitness, r_m_ (Fig 2). Significant differences in the DIC of the individual thermal performance curves translate to significant differences in overall fitness curves between populations. The key summary statistics for these models are given in SI Table 8. The lowest CTmin was 17.8°C for the Monterrey population while the highest CTmin was 19.7°C for the Acapulco population. The lowest CTmax was 31.1°C for the Jojulta population while the highest CTmax was 34.2°C for Monterrey. The lowest Topt was 26.1°C for Monterrey and the highest Topt was 28.3°C for Juárez. The population from Monterrey had both the lowest CTmin and the highest CTmax, giving it the greatest thermal breadth of 8.4°C, while the smallest Tbreadth of 6.2°C was observed for Acapulco and Jojutla populations. There were also varying Pmax values (the fitness value at the optimum temperature) with the highest Pmax being 0.504 for Cabo and the lowest Pmax being 0.358 for Jojutla. Simple correlations revealed a positive correlation between fitness Pmax and Topt (r= 0.882), as well as fitness Pmax and Tbreadth (r=0.861) (Fig 3).

**Figure 2.**
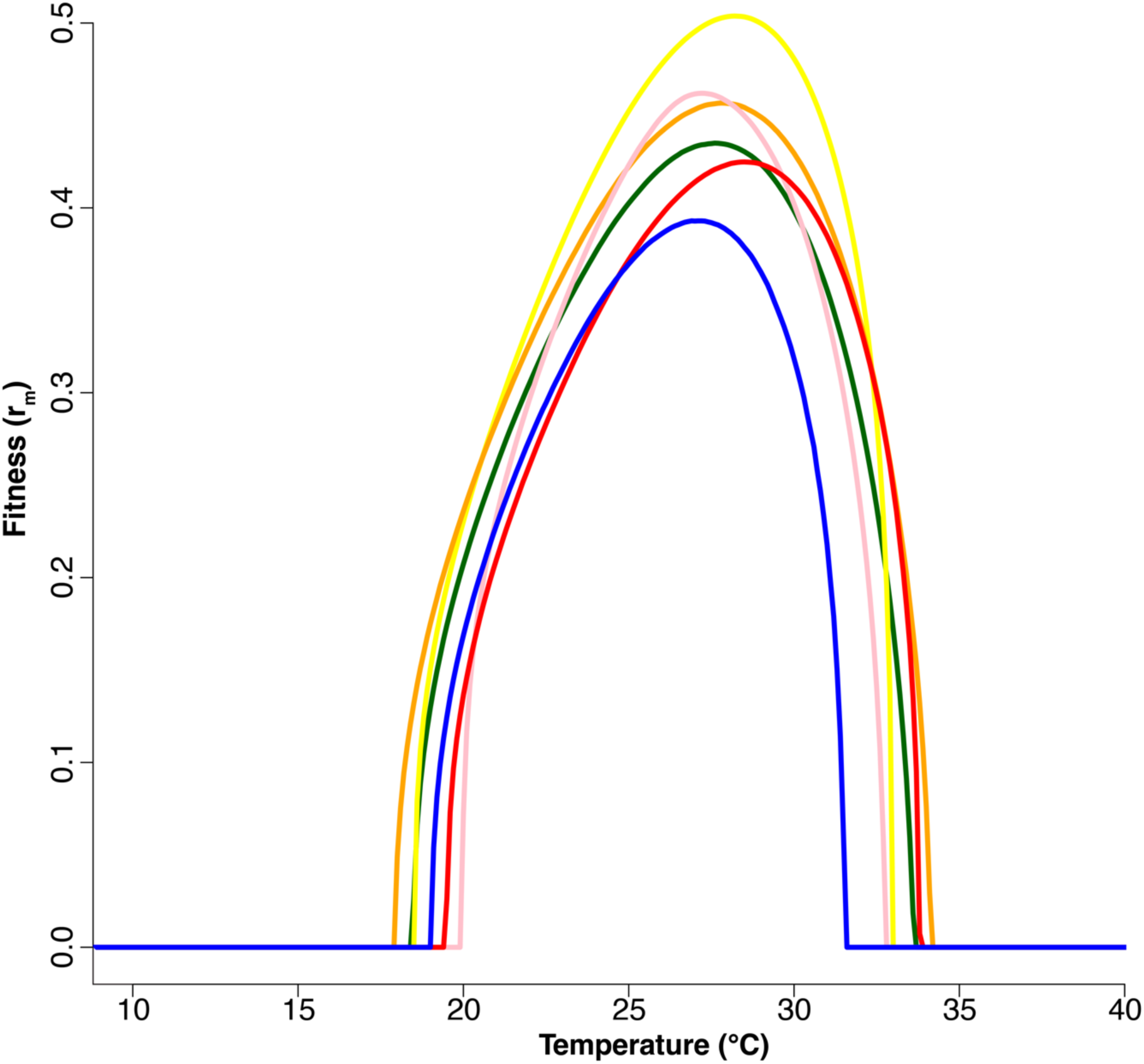
Temperature-dependent fitness (intrinsic rate of increase, r_m_) derived from individual life history traits measured for *Ae. aegypti* mosquitoes reared at 13°C, 15°C, 19°C, 23°C, 25°C, 27°C, 29°C, 31°C, 33°C, 35°C, 37°C. We measured life history traits for five populations from Mexico (Cabo San Lucas (yellow), Acapulco (pink), Monterrey (orange), Ciudad Juárez (red), and Jojutla (blue)) compared to a laboratory-adapted line (shown in green). The lines indicate mean model fits. Fits for individual populations that include the credible intervals and values are presented in SI Figure 2, SI Table 8.

**Figure 3.**
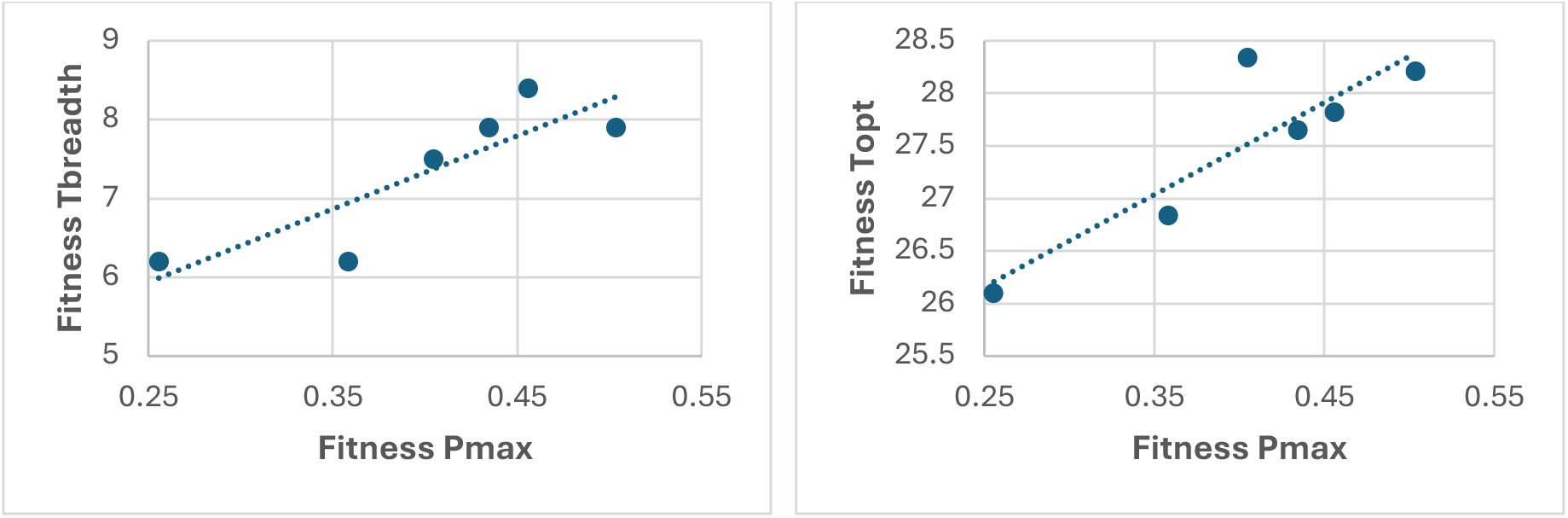
Summarized values for individual trait thermal performance curves for each population showing the relationship between (a) breadth of the curve (Tbreadth) and maximum performance (Pmax). Pearson’s correlation indicates significant strong positive relationship between Pmax and Tbreadth (r=0.882) and Pmax and Topt (r=0.861).

Finally, possible implications for transmission are presented in Fig 4, which shows the thermal performance curves for dengue virus transmission potential for each population using relative R_0_ as a comparative metric. The key summary statistics for these models are given in SI Table 9. We observe clear differences in the temperature sensitivity of transmission between individual populations. The lowest CTmin was 17.4°C for the Jojutla population and the highest CTmin was 20.2°C for the Acapulco population. The lowest CTmax was 32.8°C for Acapulco and the highest CTmax was 33.7°C for Monterrey. The lowest Topt was 26.1°C for the laboratory adapted population and the highest Topt was 28.6°C for the population from Juárez.

**Figure 4.**
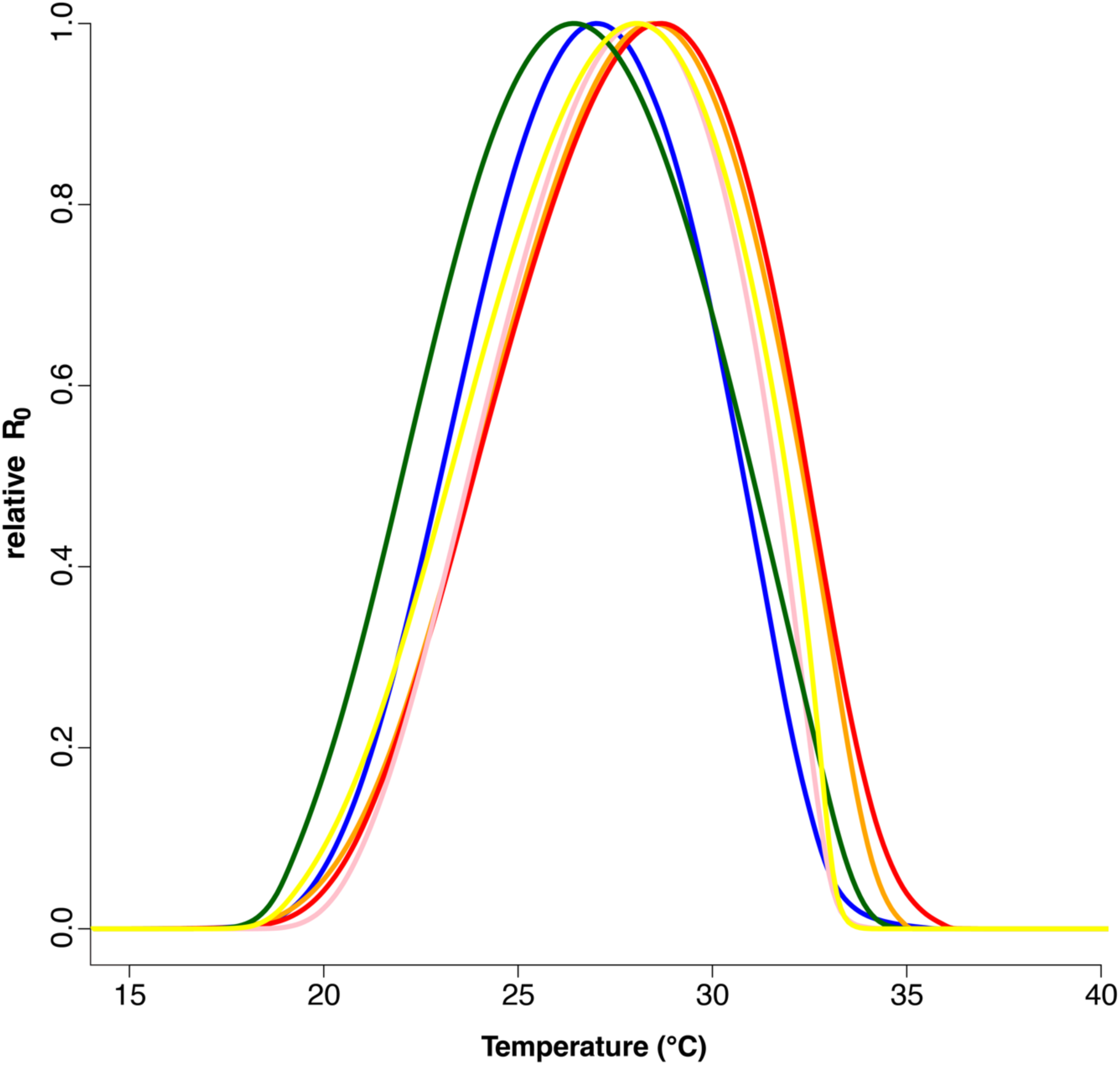
Temperature-dependent relative R_0_ (transmission rate as the basic reproduction rate R_0_) derived from individual life history traits measured for *Ae. aegypti* mosquitoes reared at 13°C, 15°C,19°C, 23°C, 25°C, 27°C, 29°C, 31°C, 33°C, 35°C, 37°C. We measured mosquito life history traits for five populations from Mexico (Cabo San Lucas (yellow), Acapulco (pink), Monterrey (orange), Ciudad Juárez (red), and Jojutla (blue)) compared to a laboratory-adapted line (shown in green). The lines indicate mean model fits. Fits for individual populations that include the credible intervals and model summary values are presented in SI Figure 3, SI Table 9.

## Discussion

In this study, we used a common-garden laboratory experiment to examine the thermal performance of five field-derived populations of *Ae. aegypti*, together with a longstanding lab strain. We examined individual life history traits as well as composite metrics for population fitness and dengue transmission potential. The results reveal significant between-population differences in several life history traits leading to distinct thermal performance curves for fitness and transmission potential. These results have important implications for understanding the effects of temperature and climate change on the transmission of vector-borne disease.

Examination of individual life history traits reveals a slightly mixed picture, with large between-population differences in thermal performance of certain traits (e.g. fecundity) but more conserved performance curves for others (e.g. larval development rate). Previous studies in a range of other taxa also reveal differences in thermal performance curves among traits and life stages (76–80), indicating that thermal responses are not necessarily co-adapted predictably across all traits (81,82). Due to this, it is predicted that the rate of thermal adaptation in an arthropod population is constrained by changes in peak performance temperatures for key life-history traits (82).

Changes in the growth rate of populations are directed by shifts in thermal performance curves of individual traits, which are thought to shift more rapidly than overall fitness with selection because of more plastic thermodynamic constraints (83,84). To determine overall differences in thermal performance between populations, we therefore combine individual life history traits into measures of population fitness. This is an important step as most studies do not characterize all traits necessary to estimate fitness (80). In this study, these fitness models further reveal multiple differences in thermal performance curves between populations (Fig 2, S2 Fig, S8 Table).

The physiological and evolutionary factors that shape variation in thermal performance curves have been the subject of much research and center around two main hypotheses. The ‘hotter-is-better’ hypothesis argues that because of thermodynamic constraints, the maximal performance of organisms with high optimal temperatures should be greater than that of organisms with low optimal temperatures and will tend to have greater thermal breadth (49,85–88). The ‘jack-of-all-temperatures’ hypothesis argues that a trade-off should exist between maximal performance and breadth of performance because different factors of a thermal performance curve can evolve independently and the association between these factors would be shown through evolutionary trade-offs (89–93). In our fitness curves, the positive correlation between maximum fitness (Pmax) and optimum temperature (Topt), as well as maximum fitness (Pmax) and thermal performance breadth (Tbreadth) (Fig 3) provide no signal of a generalist-specialist trade-off and are more consistent with the ‘hotter-is-better’ hypothesis.

The differences in thermal performance of the individual mosquito populations further lead to differences in temperature-dependent transmission potential of dengue virus (Fig 4). Interpreting these results in terms of absolute transmission risk is not possible as we did not measure infection traits and instead used previously published data for the thermal performance curves of vector competence and pathogen development rate that are not specific to our populations (10). However, as with other traits, vector competence and pathogen development rate might differ between populations (94,95). Further, estimating absolute R_0_ requires estimates of the vector-host ratio and we do not have measures of human density in the different locations, nor can we predict absolute vector densities since these could vary due to a range of factors independent of temperature. Nonetheless, the estimates of relative R_0_ enable us to draw several important insights concerning the possible effects of temperature and temperature change on dengue transmission potential with local adaptation of the mosquito vector.

First, transmission potential is expected to differ between populations at particular temperatures. For example, at 29°C the population from Ciudad Juárez is at its maximum transmission potential whereas the population from Jojutla is only at 60%. Conversely, at 25°C the lab and Jojutla populations are at their maximum, whereas Ciudad Juárez is only 60%. Further towards the thermal limits, at 20°C the transmission potential for Acapulco is close to zero yet relative R_0_ for the lab population is 15%, while at 33°C both Cabo San Lucas and Acapulco are close to zero, yet Monterrey and Ciudad Juárez have relative R_0_ values above 40%. These are substantial effect sizes and clearly demonstrate that thermal dependence of transmission potential differs between populations and is not well represented by any of the single models. To illustrate this point further, the generic thermal performance curve for relative R_0_ developed by Mordecai et al. (2017) using data from multiple published studies has a Topt of 29.1°C (95% CI: 28.4-29.8°C), a CTmin of 17.8°C (95% CI: 14.6-21.2°C) and CTmax of 34.6°C (95% CI: 34.1-35.6°C)(10). In our relative R_0_ models we see the Topt of the individual populations ranging from 26.1 (95% CI: 27.1-25.9°C) to 28.4°C (95% CI: 28.3-28.5°C), CTmin from 17.4 (95% CI: 16.6-23.0°C) to 20.2°C (95% CI:23.1-18.7°C), and CTmax from 32.7 (95% CI:31.3-33.7°C) – 36.1°C (95% CI:28.6-36.9°C). As an additional observation, while our field populations differ from one another, the largest outlier is the long-standing lab strain. Accordingly, our results provide a possible cautionary note to the many studies that explore aspects of transmission of vector-borne diseases using data from lab strains alone. This result might be expected among long-standing lab strains as they are often maintained in constant conditions over many years.

Second, a common approach to explore the possible implications of climate warming is to track changes in transmission across the R_0_ thermal performance curve (7,9–11,22,28,36–38). With a single curve, a particular shift in temperature yields a predictable response. However, if thermal response curves differ between individual populations the responses to shifts in temperature will be more idiosyncratic. For example, a warming of 3°C from 26 to 29°C would suggest an increase in relative R_0_ of around 10% for Monterrey but a decline of 30% for Jojutla. A shift from 29 to 32°C would cause a decline in transmission potential in all populations but for Ciudad Juárez the relative R_0_ would still be at 25%, whereas transmission potential for Acapulco would be almost zero.

Third, the standard approach of applying thermal performance curves to predict future changes in transmission due to warming assumes that the thermal performance curves themselves remain static. However, the fact that we see variation in thermal performance between individual populations indicates thermal performance curves are not fixed. So, while short-term changes in temperature might shift transmission along existing thermal performance curves (43), adaptation could modify the thermal responses in the medium- to longer-term (48). While absolute limits to adaptation will exist (50,96,97) ongoing adaptation to a changing environment could make responses to climate change more dynamic.

Previous research suggests genetic differentiation between discrete populations of *Ae. aegypti* in Mexico sampled across similar spatial scales to ours (98,99). However, our study examined phenotypic variation only, so we cannot say whether there are genetic differences underpinning variation in thermal performance. Furthermore, because we examined a relatively small number of mosquito populations, we have limited power to correlate observed differences in thermal performance curves with specific features of the respective home environments. Hence, it is difficult to conclude that the patterns we observe are a result of adaptation to temperature alone. Research on populations of *Drosophila* species and the mosquito *Culex tarsalis* have demonstrated correlations between thermal performance and proxies of environmental temperature such as gradients in latitude or altitude (61,62). Other research on the mosquito *Nyssorhynchus darlingi* indicates that variation in life history traits between populations results from both genetic differences among localities as well as plastic responses to differences in temperature (65). It is likely, therefore, that the phenotypic differences we observe between populations are a consequence, at least in part, of adaptive responses to temperature. Indeed, in previous research we used experimental passage to demonstrate that differences in thermal performance curves could be generated within 10 generations in response to differences in background temperature alone (48). Whether the differences are due to genetic adaptation, phenotypic plasticity, or a combination of both, does not alter the functional significance of our results.

Overall, our study demonstrates between-population differences in thermal performance *Ae. aegypti* mosquitoes that influence the predicted effects of temperature on mosquito fitness and dengue transmission potential at the local level. These results add complexities for extrapolating single thermal performance models over space and time, since existing adaptation could result in idiosyncratic responses of individual populations, and future local adaptation could further shift thermal performance curves. Given the public health significance of the multitude of pathogens transmitted by *Aedes* and other mosquitoes (e.g. dengue, Zika, Chikungunya, Yellow Fever, malaria, West Nile, filariasis, Ross River virus) and interests in the influence of climate change, this research highlights the importance of better characterizing the thermal dependence of mosquitoes and their pathogens. More generally, the interacting effects of local adaption and climate change could extend to many systems, including other vector-borne diseases of humans (e.g. Chagas disease transmitted by triatomine bugs (100–102)), animals (e.g. Bluetongue virus transmitted by culicoid midges (103–106)) and plants (e.g. numerous viruses transmitted by Hemiptera (107,108).

## Data Availability

The data that support the findings of this study will be available in Dryad.

## Permissions

All experiments were conducted under Penn State IBC protocol # 48219. Mosquito populations were imported under CDC import permit # 2018-03-181.

## Acknowledgements

We would like to thank the members of the Matthew Thomas, Elizabeth McGraw, Andrew Read, and David Kennedy lab groups for their helpful discussion and input.

## References

1. Brady OJ, Gething PW, Bhatt S, Messina JP, Brownstein JS, Hoen AG, Moyes CL, Farlow AW, Scott TW, Hay SI. Refining the global spatial limits of dengue virus transmission by evidence-based consensus. PLoS Negl Trop Dis. 2012;6(8):e1760. 10.1371/journal.pntd.000176.

2. Mayer SV, Tesh RB, Vasilakis N. The emergence of arthropod-borne viral diseases: A global prospective on dengue, chikungunya and zika fevers. Acta tropica. 2017 Feb 1;166:155–63. 10.1016/j.actatropica.2016.11.020

3. Teixeira MG, Siqueira Jr JB, Ferreira GL, Bricks L, Joint G. Epidemiological trends of dengue disease in Brazil (2000–2010): a systematic literature search and analysis. PLoS neglected tropical diseases. 2013 Dec 19;7(12):e2520. 10.1371/journal.pntd.0002520

4. Chen SC, Hsieh MH. Modeling the transmission dynamics of dengue fever: implications of temperature effects. Science of the total environment. 2012 Aug 1;431:385–91. 10.1016/j.scitotenv.2012.05.012

5. Lee H, Kim JE, Lee S, Lee CH. Potential effects of climate change on dengue transmission dynamics in Korea. PLoS One. 2018 Jun 28;13(6):e0199205. 10.1371/journal.pone.0199205

6. Thomas MB. Epidemics on the move: Climate change and infectious disease. PLoS biology. 2020 Nov 24;18(11):e3001013. 10.1371/journal.pbio.3001013

7. Ryan SJ, Carlson CJ, Mordecai EA, Johnson LR. Global expansion and redistribution of Aedes-borne virus transmission risk with climate change. PLoS neglected tropical diseases. 2019 Mar 28;13(3):e0007213. 10.1371/journal.pntd.0007213

8. Åström C, Rocklöv J, Hales S, Béguin A, Louis V, Sauerborn R. Potential distribution of dengue fever under scenarios of climate change and economic development. Ecohealth. 2012 Dec;9:448–54. 10.1007/s10393-012-0808-0

9. Ryan SJ, Carlson CJ, Tesla B, Bonds MH, Ngonghala CN, Mordecai EA, Johnson LR, Murdock CC. Warming temperatures could expose more than 1.3 billion new people to Zika virus risk by 2050. Global Change Biology. 2021 Jan;27(1):84–93. 10.1111/gcb.15384

10. Mordecai EA, Cohen JM, Evans MV, Gudapati P, Johnson LR, Lippi CA, Miazgowicz K, Murdock CC, Rohr JR, Ryan SJ, Savage V. Detecting the impact of temperature on transmission of Zika, dengue, and chikungunya using mechanistic models. PLoS neglected tropical diseases. 2017 Apr 27;11(4):e0005568. 10.1371/journal.pntd.0005568

11. Shocket MS, Verwillow AB, Numazu MG, Slamani H, Cohen JM, El Moustaid F, Rohr J, Johnson LR, Mordecai EA. Transmission of West Nile and five other temperate mosquito-borne viruses peaks at temperatures between 23 C and 26 C. Elife. 2020 Sep 15;9:e58511. 10.7554/eLife.58511

12. Liu-Helmersson J, Quam M, Wilder-Smith A, Stenlund H, Ebi K, Massad E, Rocklöv J. Climate change and Aedes vectors: 21st century projections for dengue transmission in Europe. EBioMedicine. 2016 May 1;7:267–77. 10.1016/j.ebiom.2016.03.046

13. Colón-González FJ, Sewe MO, Tompkins AM, Sjödin H, Casallas A, Rocklöv J, Caminade C, Lowe R. Projecting the risk of mosquito-borne diseases in a warmer and more populated world: a multi-model, multi-scenario intercomparison modelling study. The Lancet Planetary Health. 2021 Jul 1;5(7):e404–14. 10.1016/S2542-5196(21)00132-7

14. Romanello M, Di Napoli C, Green C, Kennard H, Lampard P, Scamman D, Walawender M, Ali Z, Ameli N, Ayeb-Karlsson S, Beggs PJ. The 2023 report of the Lancet Countdown on health and climate change: the imperative for a health-centred response in a world facing irreversible harms. The Lancet. 2023 Dec 16;402(10419):2346–94. 10.1016/S0140-6736(23)01859-7

15. Huey RB, Berrigan D. Temperature, demography, and ectotherm fitness. The American Naturalist. 2001 Aug;158(2):204–10. 10.1086/321314

16. Angilletta MJ. Thermal Adaptation: A Theoretical and Empirical Synthesis. Thermal Adaptation: A Theoretical and Empirical Synthesis. 2009. 10.1093/acprof:oso/9780198570875.001.1

17. Dell AI, Pawar S, Savage VM. Systematic variation in the temperature dependence of physiological and ecological traits. Proceedings of the National Academy of Sciences. 2011 Jun 28;108(26):10591–6. 10.1073/pnas.1015178108

18. Delatte H, Gimonneau G, Triboire A, Fontenille D. Influence of temperature on immature development, survival, longevity, fecundity, and gonotrophic cycles of Aedes albopictus, vector of chikungunya and dengue in the Indian Ocean. Journal of medical entomology. 2009 Jan 1;46(1):33–41. 10.1603/033.046.0105

19. Paaijmans KP, Read AF, Thomas MB. Understanding the link between malaria risk and climate. Proceedings of the National Academy of Sciences. 2009 Aug 18;106(33):13844–9. 10.1073/PNAS.0903423106

20. Paaijmans KP, Blanford S, Bell AS, Blanford JI, Read AF, Thomas MB. Influence of climate on malaria transmission depends on daily temperature variation. Proc Natl Acad Sci U S A. 2010 Aug 24;107(34):15135–9. 10.1073/pnas.1006422107

21. Lambrechts L, Paaijmans KP, Fansiri T, Carrington LB, Kramer LD, Thomas MB, Scott TW. Impact of daily temperature fluctuations on dengue virus transmission by Aedes aegypti. Proc Natl Acad Sci U S A. 2011 May 3;108(18):7460–5. 10.1073/pnas.1101377108

22. Mordecai EA, Paaijmans KP, Johnson LR, Balzer C, Ben-Horin T, de Moor E, et al. Optimal temperature for malaria transmission is dramatically lower than previously predicted. Ecol Lett. 2013 Jan;16(1):22–30. 10.1111/ele.12015

23. Paaijmans KP, Heinig RL, Seliga RA, Blanford JI, Blanford S, Murdock CC, Thomas MB. Temperature variation makes ectotherms more sensitive to climate change. Global change biology. 2013 Aug;19(8):2373–80. 10.1111/gcb.12240

24. Grigaltchik VS, Webb C, Seebacher F. Temperature modulates the effects of predation and competition on mosquito larvae. Ecological Entomology. 2016 Dec;41(6):668–75. 10.1111/een.12339

25. Shapiro LL, Whitehead SA, Thomas MB. Quantifying the effects of temperature on mosquito and parasite traits that determine the transmission potential of human malaria. PLoS biology. 2017 Oct 16;15(10):e2003489. 10.1371/journal.pbio.2003489

26. Villarreal SM, Winokur O, Harrington L. The impact of temperature and body size on fundamental flight tone variation in the mosquito vector Aedes aegypti (Diptera: Culicidae): implications for acoustic lures. Journal of medical entomology. 2017 Sep 1;54(5):1116–21. 10.1093/jme/tjx079

27. Reiskind MH, Janairo MS. Tracking Aedes aegypti (Diptera: Culicidae) larval behavior across development: effects of temperature and nutrients on individuals’ foraging behavior. Journal of medical entomology. 2018 Aug 29;55(5):1086–92. 10.1093/jme/tjy073

28. Tesla B, Demakovsky LR, Mordecai EA, Ryan SJ, Bonds MH, Ngonghala CN, Brindley MA, Murdock CC. Temperature drives Zika virus transmission: evidence from empirical and mathematical models. Proceedings of the Royal Society B. 2018 Aug 15;285(1884):20180795. 10.1098/rspb.2018.0795

29. Waite JL, Suh E, Lynch PA, Thomas MB. Exploring the lower thermal limits for development of the human malaria parasite, Plasmodium falciparum. Biology letters. 2019 Jun 28;15(6):20190275. 10.1098/rsbl.2019.0275

30. Suh E, Grossman MK, Waite JL, Dennington NL, Sherrard-Smith E, Churcher TS, Thomas MB. The influence of feeding behaviour and temperature on the capacity of mosquitoes to transmit malaria. Nature ecology & evolution. 2020 Jul;4(7):940–51. 10.1038/s41559-020-1182-x

31. Zouache K, Fontaine A, Vega-Rua A, Mousson L, Thiberge JM, Lourenco-De-Oliveira R, Caro V, Lambrechts L, Failloux AB. Three-way interactions between mosquito population, viral strain and temperature underlying chikungunya virus transmission potential. Proceedings of the Royal Society B: Biological Sciences. 2014 Oct 7;281(1792):20141078. 10.1098/rspb.2014.1078

32. Ruybal JE, Kramer LD, Kilpatrick AM. Geographic variation in the response of Culex pipiens life history traits to temperature. Parasites & vectors. 2016 Dec;9:1–9. 10.1186/s13071-016-1402-z

33. Shapiro LL, Murdock CC, Jacobs GR, Thomas RJ, Thomas MB. Larval food quantity affects the capacity of adult mosquitoes to transmit human malaria. Proceedings of the Royal Society B: Biological Sciences. 2016 Jul 13;283(1834):20160298. 10.1098/rspb.2016.0298

34. Huxley PJ, Murray KA, Pawar S, Cator LJ. The effect of resource limitation on the temperature dependence of mosquito population fitness. Proceedings of the Royal Society B. 2021 Apr 28;288(1949):20203217. 10.1098/rspb.2020.3217

35. Huber JH, Childs ML, Caldwell JM, Mordecai EA. Seasonal temperature variation influences climate suitability for dengue, chikungunya, and Zika transmission. PLoS neglected tropical diseases. 2018 May 10;12(5):e0006451. 10.1371/journal.pntd.0006451

36. Mordecai EA, Ryan SJ, Caldwell JM, Shah MM, LaBeaud AD. Climate change could shift disease burden from malaria to arboviruses in Africa. The Lancet Planetary Health. 2020 Sep 1;4(9):e416–23. 10.1016/S2542-5196(20)30178-9

37. Johnson LR, Ben-Horin T, Lafferty KD, McNally A, Mordecai E, Paaijmans KP, Pawar S, Ryan SJ. Understanding uncertainty in temperature effects on vector-borne disease: a Bayesian approach. Ecology. 2015 Jan;96(1):203–13. 10.1890/13-1964.1

38. Caldwell JM, LaBeaud AD, Lambin EF, Stewart-Ibarra AM, Ndenga BA, Mutuku FM, Krystosik AR, Ayala EB, Anyamba A, Borbor-Cordova MJ, Damoah R. Climate predicts geographic and temporal variation in mosquito-borne disease dynamics on two continents. Nature communications. 2021 Feb 23;12(1):1233. 10.1038/s41467-021-21496-7

39. Parham PE, Pople D, Christiansen-Jucht C, Lindsay S, Hinsley W, Michael E. Modeling the role of environmental variables on the population dynamics of the malaria vector Anopheles gambiae sensu stricto. Malaria Journal. 2012 Dec;11:1–3. 10.1186/1475-2875-11-271

40. Caminade C, Kovats S, Rocklov J, Tompkins AM, Morse AP, Colón-González FJ, Stenlund H, Martens P, Lloyd SJ. Impact of climate change on global malaria distribution. Proceedings of the National Academy of Sciences. 2014 Mar 4;111(9):3286–91. 10.1073/pnas.1302089111

41. Liu-Helmersson J, Stenlund H, Wilder-Smith A, Rocklöv J. Vectorial capacity of Aedes aegypti: effects of temperature and implications for global dengue epidemic potential. PloS one. 2014 Mar 6;9(3):e89783. 10.1371/journal.pone.0089783

43. Sternberg ED, Thomas MB. Local adaptation to temperature and the implications for vector-borne diseases. Trends in Parasitology. 2014 Mar 1;30(3):115–22. 10.1016/j.pt.2013.12.010

44. Kingsolver JG, Massie KR, Ragland GJ, Smith MH. Rapid population divergence in thermal reaction norms for an invading species: breaking the temperature–size rule. Journal of Evolutionary Biology. 2007 May 1;20(3):892–900. 10.1111/J.1420-9101.2007.01318.X

45. Deutsch CA, Tewksbury JJ, Huey RB, Sheldon KS, Ghalambor CK, Haak DC, Martin PR. Impacts of climate warming on terrestrial ectotherms across latitude. Proceedings of the National Academy of Sciences. 2008 May 6;105(18):6668–72. 10.1073/pnas.0709472105

46. Larson EL, Tinghitella RM, Taylor SA. Insect hybridization and climate change. Frontiers in Ecology and Evolution. 2019 Sep 20;7:348. 10.3389/fevo.2019.00348

47. Overgaard J, Sørensen JG. Rapid thermal adaptation during field temperature variations in Drosophila melanogaster. Cryobiology. 2008 Apr 1;56(2):159–62. 10.1016/j.cryobiol.2008.01.001

48. Dennington NL, Grossman MK, Ware-Gilmore F, Teeple JL, Johnson LR, Shocket MS, McGraw EA, Thomas MB. Phenotypic adaptation to temperature in the mosquito vector, Aedes aegypti. Global Change Biology. 2024 Jan;30(1):e17041. 10.1111/gcb.17041

49. Alruiz JM, Peralta-Maraver I, Bozinovic F, Santos M, Rezende EL. Temperature adaptation and its impact on the shape of performance curves in Drosophila populations. Proceedings of the Royal Society B. 2023 May 10;290(1998):20230507. 10.1098/rspb.2023.0507

50. Kellermann V, Van Heerwaarden B, Sgrò CM, Hoffmann AA. Fundamental evolutionary limits in ecological traits drive Drosophila species distributions. Science. 2009 Sep 4;325(5945):1244–6. 10.1126/science.1175443

51. Bennett JM, Sunday J, Calosi P, Villalobos F, Martínez B, Molina-Venegas R, Araújo MB, Algar AC, Clusella-Trullas S, Hawkins BA, Keith SA. The evolution of critical thermal limits of life on Earth. Nature communications. 2021 Feb 19;12(1):1198. 10.1038/s41467-021-21263-8

52. Weaving H, Terblanche JS, Pottier P, English S. Meta-analysis reveals weak but pervasive plasticity in insect thermal limits. Nature communications. 2022 Sep 8;13(1):5292. 10.1038/s41467-022-32953-2

53. Johansson J. Evolutionary responses to environmental changes: how does competition affect adaptation?. Evolution. 2008 Feb 1;62(2):421–35. 10.1111/j.1558-5646.2007.00301.x

54. Bürger R, Lynch M. Evolution and extinction in a changing environment: a quantitative-genetic analysis. Evolution. 1995 Feb;49(1):151–63. 10.1111/j.1558-5646.1995.tb05967.x

55. Lynch M, Lande R. Evolution and extinction in response to environmental change. Biotic Interactions and Global Change. 1993;

56. Orr HA, Unckless RL. Population extinction and the genetics of adaptation. The American Naturalist. 2008 Aug;172(2):160–9. 10.1086/589460

57. Urbanski J, Mogi M, O’Donnell D, DeCotiis M, Toma T, Armbruster P. Rapid adaptive evolution of photoperiodic response during invasion and range expansion across a climatic gradient. The American Naturalist. 2012 Apr 1;179(4):490–500. 10.1086/664709

58. Gatton ML, Chitnis N, Churcher T, Donnelly MJ, Ghani AC, Godfray HC, Gould F, Hastings I, Marshall J, Ranson H, Rowland M. The importance of mosquito behavioural adaptations to malaria control in Africa. Evolution. 2013 Apr 1;67(4):1218–30. 10.1111/evo.12063

59. Egizi A, Fefferman NH, Fonseca DM. Evidence that implicit assumptions of ‘no evolution’of disease vectors in changing environments can be violated on a rapid timescale. Philosophical Transactions of the Royal Society B: Biological Sciences. 2015 Apr 5;370(1665):20140136. 10.1098/rstb.2014.0136

60. Ranson H, Lissenden N. Insecticide resistance in African Anopheles mosquitoes: a worsening situation that needs urgent action to maintain malaria control. Trends in parasitology. 2016 Mar 1;32(3):187–96. 10.1016/j.pt.2015.11.010

61. Sgrò CM, Overgaard J, Kristensen TN, Mitchell KA, Cockerell FE, Hoffmann AA. A comprehensive assessment of geographic variation in heat tolerance and hardening capacity in populations of Drosophila melanogaster from eastern Australia. Journal of evolutionary biology. 2010 Nov 1;23(11):2484–93. 10.1111/j.1420-9101.2010.02110.x

62. Vorhees AS, Gray EM, Bradley TJ. Thermal resistance and performance correlate with climate in populations of a widespread mosquito. Physiological and Biochemical Zoology. 2013 Jan 1;86(1):73–81. 10.1086/668851

63. Merilä J, Hendry AP. Climate change, adaptation, and phenotypic plasticity: the problem and the evidence. Evolutionary applications. 2014 Jan;7(1):1–4. 10.1111/eva.12137

64. de Villemereuil P, Gaggiotti OE, Mouterde M, Till-Bottraud I. Common garden experiments in the genomic era: new perspectives and opportunities. Heredity. 2016 Mar;116(3):249–54. 10.1038/hdy.2015.93

65. Chu VM, Sallum MA, Moore TE, Lainhart W, Schlichting CD, Conn JE. Regional variation in life history traits and plastic responses to temperature of the major malaria vector Nyssorhynchus darlingi in Brazil. Scientific reports. 2019 Mar 29;9(1):5356. 10.1038/s41598-019-41651-x

66. Gachohi JM, Njenga MK, Kitala P, Bett B. Modelling Vaccination Strategies against Rift Valley Fever in Livestock in Kenya. PLoS Negl Trop Dis. 2016 Dec 14;10(12) e0005049. 10.1371/journal.pntd.0005049

67. Harrington LC, Fleisher A, Ruiz-Moreno D, Vermeylen F, Wa C V., Poulson RL, et al. Heterogeneous Feeding Patterns of the Dengue Vector, Aedes aegypti, on Individual Human Hosts in Rural Thailand. PLoS Negl Trop Dis. 2014 Aug 7;8(8): e3048. 10.1371/journal.pntd.0003048

68. Liao CM, Huang TL, Cheng YH, Chen WY, Hsieh NH, Chen SC, et al. Assessing dengue infection risk in the southern region of Taiwan: Implications for control. Epidemiol Infect. 2015 Apr;143(5): 1059–72. 10.1017/S0950268814001745

69. Index R, Development TR, Team C. The R Environment for Statistical Computing and Graphics. Vol. 1, Development. 2003.

70. Plummer M, Best N, Cowles K, Vines K. CODA: convergence diagnosis and output analysis for MCMC. R news. 2006 Mar;6(1):7–11.

71. Su YS, Yajima M. R2jags: a package for running jags from R. R package version 0.03-08.

72. Plummer M, Best N, Cowles K, Vines K. {CODA}: Convergence Diagnosis and Output Analysis for {MCMC}. R News. 2006;6(1).

73. Amarasekare P, Savage V. A framework for elucidating the temperature dependence of fitness. The American Naturalist. 2012 Feb 1;179(2):178–91. 10.1086/663677

74. Angilletta Jr MJ, Niewiarowski PH, Navas CA. The evolution of thermal physiology in ectotherms. Journal of thermal Biology. 2002 Aug 1;27(4):249–68. 10.1016/S0306-4565(01)00094-8

75. MacLean HJ, Sørensen JG, Kristensen TN, Loeschcke V, Beedholm K, Kellermann V, Overgaard J. Evolution and plasticity of thermal performance: an analysis of variation in thermal tolerance and fitness in 22 Drosophila species. Philosophical Transactions of the Royal Society B. 2019 Aug 5;374(1778):20180548. 10.1098/rstb.2018.0548

76. Kingsolver JG, Woods HA. Thermal sensitivity of growth and feeding in Manduca sexta caterpillars. Physiological Zoology. 1997 Nov;70(6):631–8. 10.1086/515872

77. Stevenson RD, Peterson CR, Tsuji JS. The thermal dependence of locomotion, tongue flicking, digestion, and oxygen consumption in the wandering garter snake. Physiological Zoology. 1985 Jan 1;58(1):46–57. 10.1086/physzool.58.1.30161219

78. Woods HA, Harrison JF. Interpreting rejections of the beneficial acclimation hypothesis: when is physiological plasticity adaptive?. Evolution. 2002 Sep;56(9):1863–6. 10.1111/j.0014-3820.2002.tb00201.x

79. David JR, Gibert P, Legout H, Pétavy G, Capy P, Moreteau B. Isofemale lines in Drosophila: an empirical approach to quantitative trait analysis in natural populations. Heredity. 2005 Jan;94(1):3–12. 10.1038/sj.hdy.6800562

80. Kellermann V, Chown SL, Schou MF, Aitkenhead I, Janion-Scheepers C, Clemson A, Scott MT, Sgrò CM. Comparing thermal performance curves across traits: how consistent are they?. Journal of Experimental Biology. 2019 Jun 1;222(11):jeb193433. 10.1242/jeb.193433

81. Angilletta Jr MJ, Bennett AF, Guderley H, Navas CA, Seebacher F, Wilson RS. Coadaptation: a unifying principle in evolutionary thermal biology. Physiological and Biochemical Zoology. 2006 Mar;79(2):282–94. 10.1086/499990

82. Pawar S, Huxley PJ, Smallwood TR, Nesbit ML, Chan AH, Shocket MS, Johnson LR, Kontopoulos DG, Cator LJ. Variation in temperature of peak trait performance constrains adaptation of arthropod populations to climatic warming. Nature Ecology & Evolution. 2024 Jan 25:1–1. 10.1038/s41559-023-02301-8

83. Kingsolver JG. The Well-Temperatured Biologist: (American Society of Naturalists Presidential Address). The American Naturalist. 2009 Dec;174(6):755–68. 10.1086/648310

84. Kontopoulos DG, van Sebille E, Lange M, Yvon-Durocher G, Barraclough TG, Pawar S. Phytoplankton thermal responses adapt in the absence of hard thermodynamic constraints. Evolution. 2020 Apr 1;74(4):775–90. 10.1111/evo.13946

85. Bennett AF. Evolution of the control of body temperature: is warmer better. Comparative physiology: life in water and on land. 1987 Jul 27;9:421–31.

86. Huey RB, Kingsolver JG. Evolution of thermal sensitivity of ectotherm performance. Trends in ecology & evolution. 1989 May 1;4(5):131–5. 10.1016/0169-5347(89)90211-5

87. Savage VM, Gillooly JF, Brown JH, West GB, Charnov EL. Effects of body size and temperature on population growth. The American Naturalist. 2004 Mar;163(3):429–41. 10.1086/381872

88. Santos M. Evolution of total net fitness in thermal lines: Drosophila subobscura likes it ‘warm’. Journal of Evolutionary Biology. 2007 Nov 1;20(6):2361–70. 10.1111/j.14209101.2007.01408.x

89. Montagnes DJ, Wang Q, Lyu Z, Shao C. Evaluating thermal performance of closely related taxa: Support for hotter is not better, but for unexpected reasons. Ecological Monographs. 2022 Aug;92(3):e1517. 10.1002/ecm.1517

90. Somero GN, Hochachka PW. Biochemical adaptation to the environment. American Zoologist. 1971 Feb 1;11(1):159–67. 10.1093/icb/11.1.159

91. Izem R, Kingsolver JG. Variation in continuous reaction norms: quantifying directions of biological interest. The American Naturalist. 2005 Aug;166(2):277–89. 10.1086/431314

92. Gilchrist GW. A quantitative genetic analysis of thermal sensitivity in the locomotor performance curve of Aphidius ervi. Evolution. 1996 Aug 1;50(4):1560–72. 10.1111/j.15585646.1996.tb03928.x

93. Huey RB, Hertz PE. Is a jack-of-all-temperatures a master of none?. Evolution. 1984 Mar 1;38(2):441–4. 10.2307/2408502

94. Fansiri T, Pongsiri A, Klungthong C, Ponlawat A, Thaisomboonsuk B, Jarman RG, Scott TW, Lambrechts L. No evidence for local adaptation of dengue viruses to mosquito vector populations in T hailand. Evolutionary applications. 2016 Apr;9(4):608–18. 10.1111/eva.12360

95. Lambrechts L, Chevillon C, Albright RG, Thaisomboonsuk B, Richardson JH, Jarman RG, Scott TW. Genetic specificity and potential for local adaptation between dengue viruses and mosquito vectors. BMC evolutionary biology. 2009 Dec;9:1–1. 10.1186/1471-2148-9-160

96. Hoffmann AA, Hallas RJ, Dean JA, Schiffer M. Low potential for climatic stress adaptation in a rainforest Drosophila species. Science. 2003 Jul 4;301(5629):100–2. 10.1126/science.1084296

97. Kinzner MC, Gamisch A, Hoffmann AA, Seifert B, Haider M, Arthofer W, Schlick-Steiner BC, Steiner FM. Major range loss predicted from lack of heat adaptability in an alpine Drosophila species. Science of the Total Environment. 2019 Dec 10;695:133753. 10.1016/j.scitotenv.2019.133753

98. Gloria-Soria A, Ayala D, Bheecarry A, Calderon-Arguedas O, Chadee DD, Chiappero M, Coetzee M, Elahee KB, Fernandez-Salas I, Kamal HA, Kamgang B. Global genetic diversity of Aedes aegypti. Molecular ecology. 2016 Nov;25(21):5377–95. 10.1111/mec.13866

99. Gorrochotegui-Escalante N, Munoz MD, Fernandez-Salas I, Beaty BJ, Black 4th WC. Genetic isolation by distance among Aedes aegypti populations along the northeastern coast of Mexico. The American journal of tropical medicine and hygiene. 2000 Feb;62(2):200–9. 10.4269/AJTMH.2000.62.200

100. Carmona-Castro O, Moo-Llanes DA, Ramsey JM. Impact of climate change on vector transmission of Trypanosoma cruzi (Chagas, 1909) in North America. Medical and veterinary entomology. 2018 Mar;32(1):84–101. 10.1111/mve.12269

101. Garza M, Feria Arroyo TP, Casillas EA, Sanchez-Cordero V, Rivaldi CL, Sarkar S. Projected future distributions of vectors of Trypanosoma cruzi in North America under climate change scenarios. PLoS neglected tropical diseases. 2014 May 15;8(5):e2818. 10.1371/journal.pntd.0002818

102. Ceccarelli S, Rabinovich JE. Global climate change effects on Venezuela’s vulnerability to chagas disease is linked to the geographic distribution of five triatomine species. Journal of Medical Entomology. 2015 Nov 1;52(6):1333–43. 10.1093/jme/tjv119

103. Nicolas G, Tisseuil C, Conte A, Allepuz A, Pioz M, Lancelot R, Gilbert M. Environmental heterogeneity and variations in the velocity of bluetongue virus spread in six European epidemics. Preventive veterinary medicine. 2018 Jan 1;149:1–9. 10.1016/j.prevetmed.2017.11.005

104. Brand SP, Keeling MJ. The impact of temperature changes on vector-borne disease transmission: Culicoides midges and bluetongue virus. Journal of the Royal Society Interface. 2017 Mar31;14(128):20160481. 10.1098/rsif.2016.0481

105. Samy AM, Peterson AT. Climate change influences on the global potential distribution of bluetongue virus. PloS one. 2016 Mar 9;11(3):e0150489. 10.1371/journal.pone.0150489

106. Purse BV, Mellor PS, Rogers DJ, Samuel AR, Mertens PP, Baylis M. Climate change and the recent emergence of bluetongue in Europe. Nature reviews microbiology. 2005 Feb;3(2):171–81. 10.1038/nrmicro1090

107. Chen J, Jiang K, Wang S, Li Y, Zhang Y, Tang Z, Bu W. Climate change impacts on the potential worldwide distribution of the soybean pest, Piezodorus guildinii (Hemiptera: Pentatomidae). Journal of Economic Entomology. 2023 Jun 1;116(3):761–70. 10.1093/jee/toad058

108. Skendžić S, Zovko M, Živković IP, Lešić V, Lemić D. The impact of climate change on agricultural insect pests. Insects. 2021 May 12;12(5):440. 10.3390/insects12050440

