## Supplemental Information for "Local adaptation of the mosquito vector, *Aedes aegypti,* and implications for predicting the effects of temperature and climate change on dengue transmission"

### Supporting Information Text

**Mathematical Models.**

The full model for the i[th] observed fecundity, $F_i$, including priors, is as follows:

$$f(T) = \begin{cases} -c\,(T - T_0)(T - T_m) & \text{if } T_0 \le T \le T_m \\ 0 & \text{otherwise} \end{cases}$$

$$F_i \sim T\mathcal{N}(\mu = f(T_i), \sigma^2, 0, \infty)$$

$$c \sim U(0, 1)$$

$$T_0 \sim U(0, 24)$$

$$T_m \sim U(25, 45)$$

$$\sigma \sim U(0, 1000)$$

The full model for the i[th] observed mosquito development rate, $D_i$, including priors, is as follows:

$$f(T) = \begin{cases} -cT\,(T - T_0)(\sqrt{T_m - T}) & \text{if } T_0 \le T \le T_m \\ 0 & \text{otherwise} \end{cases}$$

$$D_i \sim T\mathcal{N}(\mu = f(T_i), \sigma^2, 0, \infty)$$

$$c \sim U(0, 1)$$

$$T_0 \sim U(0, 24)$$

$$T_m \sim U(25, 45)$$

$$\sigma \sim U(0, 1000)$$

The full model for the i[th] observed egg to adult survival probability, $Y_i$, including priors, is as follows. The quadratic function is defined as

$$f(T) = \begin{cases} -c\,(T - T_0)(T - T_m) & \text{if } T_0 \le T \le T_m \\ 0 & \text{otherwise.} \end{cases}$$

This is used to define a probability between 0 and one, such that

$$p(T_i) = f(T_i)I[f(T_i) < 1] + I[f(T_i) > 1],$$

where $I[\cdot]$ is an indicator function such that is one if the argument is true, and zero otherwise. Thus the observed data ($Y_i$ = total survived and $n_i$ = number observed) is described by a binomial distribution

$$Y_i \sim Bin(p = f(T_i), n_i)$$

and priors for parameters are

$$T_0 \sim U(0, 24)$$

$$T_m \sim U(25, 45)$$

$$c \sim exp(1).$$

For the observed survival times for adult mosquitoes, we chose prior distributions to enforce positivity of the shape parameter and of the thermal performance parameters, but that are relatively uninformative, specifically,

$k, a, b \sim$ Exp$(0.001)$.

To analyze the thermal performance curves for juvenile mortality rate($m_j$), we fit symmetric thermal response functions to our data using a Bayesian approach. Specifically, we assumed a normal likelihood distribution with temperature-dependent means specified by a quadratic equation and a constant standard deviation:

$$mj(T) = aT_i^2 + bT_i + cf(T_i)$$

$$a \sim U(0, 1)$$

$$b \sim U(-1, 0)$$

$$c \sim U(0,1)$$

When $m_j(T_i) < 0$ then we set the thermal response function to 0 to prevent negative trait values.

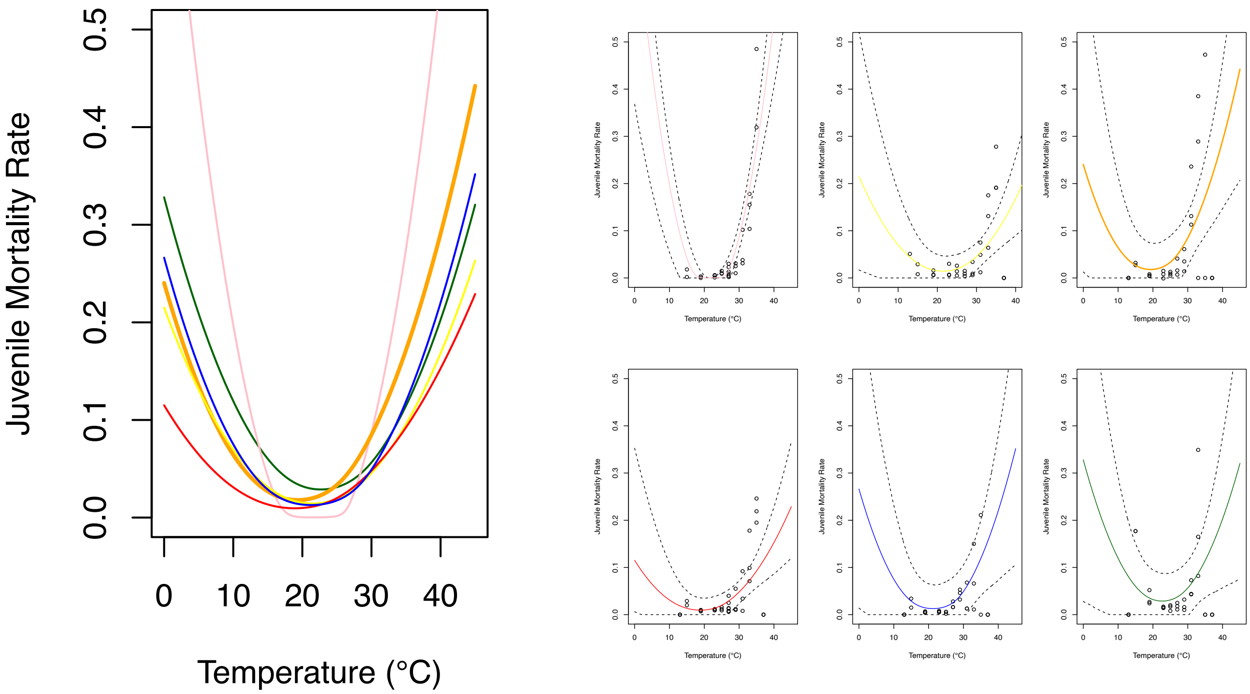


**S1 Fig**. **Juvenile mortality rate thermal performance curve.** Thermal responses for *Aedes aegypti* juvenile mortality rate for mosquitoes from Mexico compared to a laboratory line. Uninformative priors were used, and the models were fit to raw data. We measured life history traits for five populations from Mexico (Cabo San Lucas (yellow), Acapulco (pink), Monterrey (orange), Ciudad Juárez (red), and Jojutla (blue)) compared to a laboratory adapted line (shown in green). Each line was tested between 13°C-37°C, but individuals could not survive and reproduce at 13°C and 37°C.

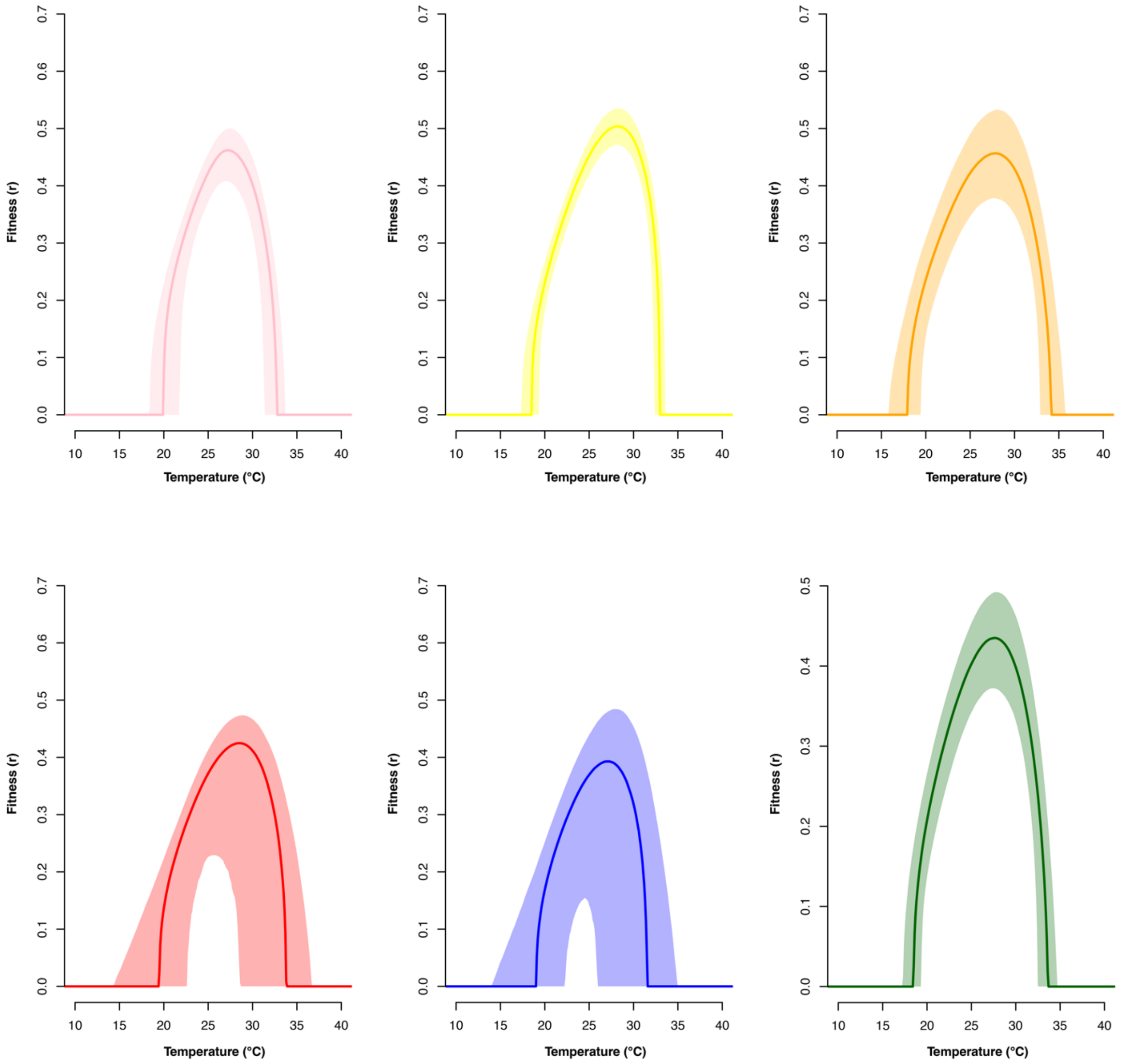


**S2 Fig. Temperature-dependent fitness models.** Temperature-dependent fitness (intrinsic rate of increase, $r_m$) derived from individual life history traits measured for *Ae. aegypti* mosquitoes reared at 13°C, 15°C, 19°C, 23°C, 25°C, 27°C, 29°C, 31°C, 33°C, 35°C, 37°C. We measured life history traits for five populations from Mexico (Cabo San Lucas (yellow), Acapulco (pink), Monterrey (orange), Ciudad Juárez (red), and Jojutla (blue)) compared to a laboratory-adapted line (shown in green). The lines indicate mean model fits while the shaded area indicates the 95% credible intervals.

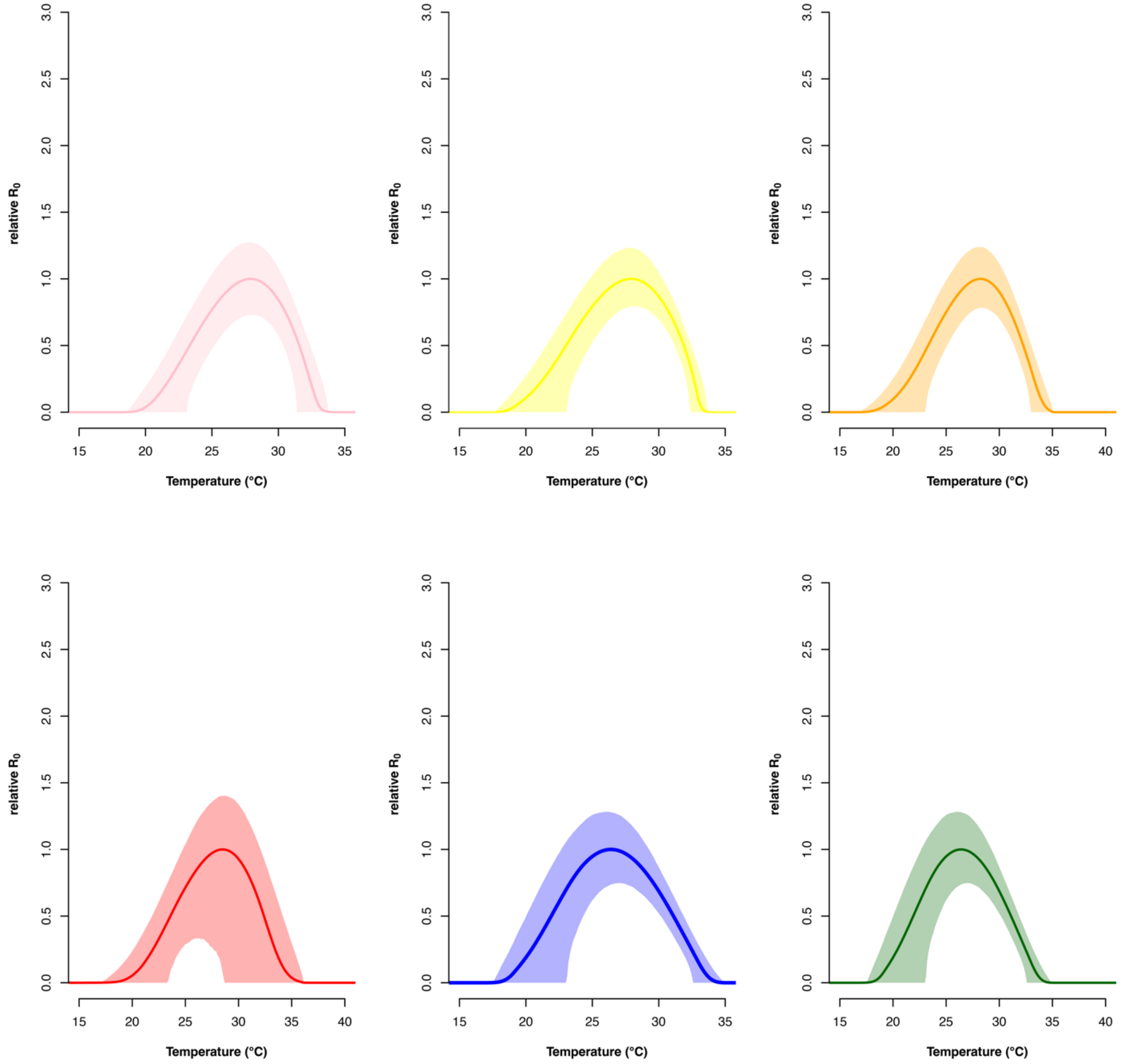


**S3 Fig. Temperature-dependent relative $R_0$ models.** Temperature-dependent relative $R_0$ (transmission rate as the basic reproduction rate $R_0$) derived from individual life history traits measured for *Ae. aegypti* mosquitoes reared at 13°C, 15°C,19°C, 23°C, 25°C, 27°C, 29°C, 31°C, 33°C, 35°C, 37°C. We measured mosquito life history traits for five populations from Mexico (Cabo San Lucas (yellow), Acapulco (pink), Monterrey (orange), Ciudad Juárez (red), and Jojutla (blue)) compared to a laboratory-adapted line (shown in green). The lines indicate mean model fits while the shaded area indicates 95% credible intervals.

**Supplementary Table 1:** Information on locations for *Aedes aegypti* mosquito collection in Mexico

| City | Elevation (m) | Latitude | Mean annual temperature (°C) | Mean annual high temperature (°C) | Mean annual low temperature (°C) | Generation from field |
|---|---|---|---|---|---|---|
| Acapulco, Guerrero | 45 | 16.85 | 27.9 | 31.3 | 23 | 2 |
| Cabo San Lucas | 10 | 22.89 | 23 | 29.9 | 18 | 3 |
| Monterrey, Nuevo Leon | 495 | 25.67 | 22.2 | 28.4 | 17 | 2 |
| Juárez, Chihuahua | 1135 | 31.69 | 17.9 | 25.8 | 10 | 3 |
| Jojutla, Morelos | 891 | 18.6 | 24.9 | 33.4 | 16.3 | 3 |

**Table S1:** Data collected over multiple years in each city. Each mean represents the grand mean across multiple years. These populations were compared to a standard laboratory adapted line (Rockefeller strain) maintained at Penn State at standard insectary conditions.

**Supplementary Table 2:** Briere model output for mosquito development rate for *Aedes aegypti* from Mexico compared to a laboratory adapted line.

| Population | $T_0$ | | | $T_m$ | | | *c* | | | DIC |
|---|---|---|---|---|---|---|---|---|---|---|
| | Mean | 95% CI | | Mean | 95% CI | | Mean | 95% CI | | |
| Acapulco, Guerrero | 11.08 | 8.847 | 12.85 | 36.814 | 36.399 | 37.018 | $9.373^{-05}$ | $8.065^{-05}$ | $1.072^{-04}$ | -201.2033 |
| Cabo San Lucas | 11.418 | 10.210 | 12.499 | 36.935 | 36.754 | 37.011 | $9.649^{-05}$ | $8.905^{-05}$ | $1.039^{-04}$ | -214.3716 |
| Monterrey, Nuevo Leon | 9.787 | 2.448 | 14.311 | 36.102 | 35.338 | 37.002 | $8.654^{-05}$ | $5.567^{-05}$ | $1.205^{-04}$ | -128.3117 |
| Juárez, Chihuahua | 11.329 | 10.239 | 12.377 | 36.938 | 36.756 | 37.008 | $9.076^{-05}$ | $8.413^{-05}$ | $9.778^{-05}$ | -241.3215 |
| Jojutla, Morelos | 11.227 | 7.872 | 13.812 | 36.346 | 35.690 | 36.991 | $9.605^{-05}$ | $7.519^{-05}$ | $1.175^{-04}$ | -162.6717 |
| Laboratory Population | 11.041 | 7.963 | 13.331 | 35.919 | 35.317 | 36.758 | $9.202^{-05}$ | $7.223^{-05}$ | $1.114^{-04}$ | -165.2648 |

| Population | Pmax | | | Topt | | | Breadth | | |
|---|---|---|---|---|---|---|---|---|---|
| | Mean | 95% CI | | Mean | 95% CI | | Mean | 95% CI | |
| Acapulco, Guerrero | 0.1391692 | 0.1316232 | 0.1465687 | 30.8 | 30.7 | 30.8 | 9.7 | 9.5 | 9.8 |
| Cabo San Lucas | 0.1424445 | 0.1373978 | 0.147592 | 31.0 | 31.0 | 31.1 | 9.6 | 9.5 | 9.6 |
| Monterrey, Nuevo Leon | 0.1262943 | 0.1090853 | 0.1433593 | 30.1 | 30.1 | 30.2 | 9.6 | 9.3 | 9.8 |
| Juárez, Chihuahua | 0.1346993 | 0.130463 | 0.1390729 | 30.9 | 30.9 | 31.1 | 9.6 | 9.5 | 9.6 |
| Jojutla, Morelos | 0.1353633 | 0.1239167 | 0.1470923 | 30.4 | 30.4 | 30.4 | 9.4 | 9.2 | 9.7 |

| | | | | | | | | | |
|---|---|---|---|---|---|---|---|---|---|
| Laboratory Population | 0.1259861 | 0.115634 | 0.1365272 | 30.1 | 29.9 | 30.1 | 9.3 | 9.1 | 9.7 |

**Table S2:** Data for mosquito development rate for five populations of *Ae. aegypti* mosquitoes from Mexico compared to one laboratory adapted line were used to generate briere model. Mean and 95% credible interval (95% HPD interval) for the critical thermal minimum ($T_0$), maximum, ($T_m$), and a rate constant (*c*) are given for mosquito development rate. The Deviance Criterion Information (DIC) is given to compare model fits. We are comparing this model to one that fits all of the points together giving a DIC of -1069.219.

**Supplementary Table 3:** Quadratic model output for egg to adult survival for *Aedes aegypti* from Mexico compared to a laboratory adapted line.

| Population | $T_0$ | | | $T_m$ | | | *c* | | | DIC |
|---|---|---|---|---|---|---|---|---|---|---|
| | Mean | 95% CI | | Mean | 95% CI | | Mean | 95% CI | | |
| Acapulco, Guerrero | 12.98 | 12.89 | 13.03 | 35.86 | 37.23 | 37.64 | 7.34e-03 | 7.20e-03 | 7.47e-03 | 970.1 |
| Cabo San Lucas | 12.44 | 12.202740 | 12.65 | 37.11 | 37.00 | 37.30 | 6.35e-03 | 6.17e-03 | 6.50e-03 | 1176 |
| Monterrey, Nuevo Leon | 12.98 | 12.90 | 13.03 | 35.03 | 35.01 | 35.06 | 7.633e-03 | 7.51e-03 | 7.74e-03 | 852 |
| Juárez, Chihuahua | 11.97 | 11.72 | 12.22 | 36.98 | 36.90 | 37.04 | 6.074e-03 | 5.93e-03 | 6.22 e-03 | 1095 |
| Jojutla, Morelos | 12.17 | 11.92 | 12.4 | 35.82 | 35.64 | 36.00 | 7.002e-03 | 6.78e-03 | 7.24e-03 | 1320 |
| Laboratory Population | 13.64 | 13.42 | 13.84 | 35.83 | 35.66 | 36.01 | 7.618e-03 | 7.37e-03 | 7.86e-03 | 713.8 |

| Population | Pmax | | | Topt | | | Breadth | | |
|---|---|---|---|---|---|---|---|---|---|
| | Mean | 95% CI | | Mean | 95% CI | | Mean | 95% CI | |
| Acapulco, Guerrero | 0.9597276 | 0.9514193 | 0.9675379 | 24.4 | 24.5 | 24.4 | 10.1 | 10.1 | 10.2 |
| Cabo San Lucas | 0.9359657 | 0.9258766 | 0.945268 | 24.9 | 25.0 | 24.9 | 10.6 | 10.5 | 10.6 |
| Monterrey, Nuevo Leon | 0.9276821 | 0.9146845 | 0.9400959 | 24.0 | 24.0 | 24.0 | 9.8 | 9.8 | 9.8 |
| Juárez, Chihuahua | 0.9285334 | 0.9181776 | 0.9383473 | 24.5 | 24.6 | 24.5 | 10.3 | 10.1 | 10.3 |
| Jojutla, Morelos | 0.9666728 | 0.9582457 | 0.9747009 | 24.3 | 24.4 | 24.3 | 10.1 | 10.0 | 10.2 |
| Laboratory Population | 0.899282 | 0.8865129 | 0.9115908 | 25.0 | 25.0 | 25.0 | 8.8 | 8.8 | 8.8 |

**Table S3:** Data for egg to adult survival on *Ae. aegypti* from 5 population from Mexico compared to one laboratory adapted line were used to generate quadratic model. Mean and 95% credible interval (95% HPD interval) for the critical thermal minimum ($T_0$), maximum, ($T_m$), and a rate constant (*c*) are given for egg-to-adult survival. The Deviance Information Criterion (DIC) is given to compare model fits. We are comparing this model to one that fits all of the points together giving a DIC of 6133.

**Supplementary Table 4:** Quadratic model output for eggs per female per gonotrophic cycle for *Aedes aegypti* from Mexico compared to a laboratory adapted line.

| Population | $T_0$ | | | $T_m$ | | | *c* | | | DIC |
|---|---|---|---|---|---|---|---|---|---|---|
| | Mean | 95% CI | | Mean | 95% CI | | Mean | 95% CI | | |
| Acapulco, Guerrero | 19.991 | 18.354 | 21.790 | 32.698 | 31.338 | 33.717 | 0.830 | 0.539 | 0.993 | 2019.46 |
| Cabo San Lucas | 18.477 | 17.344 | 19.434 | 33.016 | 32.373 | 33.630 | 0.938 | 0.790 | 0999 | 1655.237 |
| Monterrey, Nuevo Leon | 17.857 | 15.857 | 19.444 | 34.235 | 32.818 | 36.145 | 0.843 | 0.554 | 0.994 | 1652.761 |
| Juárez, Chihuahua | 19.086 | 12.741 | 22.313 | 33.848 | 29.066 | 37.938 | 0.538 | 0.099 | 0.949 | 2000.809 |
| Jojutla, Morelos | 18.327 | 8.709 | 22.028 | 31.170 | 25.889 | 36.417 | 0.569 | 0.031 | 0.971 | 1671.899 |
| Laboratory Population | 18.400 | 17.259 | 19.297 | 33.706 | 32.550 | 34.856 | 0.954 | 0.833 | 0.999 | 1671.337 |

| Population | Pmax | | | Topt | | | Breadth | | |
|---|---|---|---|---|---|---|---|---|---|
| | Mean | 95% CI | | Mean | 95% CI | | Mean | 95% CI | |
| Acapulco, Guerrero | 32.79473 | 20.45213 | 41.86363 | 26.4 | 26.5 | 26.2 | 5.5 | 4.3 | 6.3 |
| Cabo San Lucas | 49.35614 | 42.54272 | 55.32652 | 25.7 | 25.8 | 25.5 | 6.4 | 5.7 | 7 |
| Monterrey, Nuevo Leon | 55.41098 | 45.28123 | 63.23186 | 26.1 | 26.1 | 25.7 | 7.2 | 6.1 | 8.4 |
| Juárez, Chihuahua | 26.09382 | 4.24679 | 39.06503 | 26.6 | 24.8 | 26.7 | 6.1 | 2.3 | 7.6 |
| Jojutla, Morelos | 19.99891 | 1.324288 | 37.01411 | 25.0 | 24.2 | 25.6 | 5.3 | 1.6 | 6.6 |
| Laboratory Population | 56.05282 | 45.55214 | 64.38806 | 26.1 | 25.8 | 26.0 | 6.7 | 5.9 | 8.0 |

**Table S4:** Data for eggs per female per day on *Ae. aegypti* mosquitoes selected over ten generations were used to generate quadratic model. Mean and 95% credible interval (95% HPD interval) for the critical thermal minimum ($T_0$), maximum, ($T_m$), and a rate constant (*c*) are given for fecundity. The Deviance Criterion Information (DIC) is given to compare model fits. We are comparing this model to one that fits all of the points together giving a DIC of 9027.19.

**Supplementary Table 5:** Weibull survival model output for adult survival.

| Population | $y_i$ | | | β | | | $k$ | | | DIC |
|---|---|---|---|---|---|---|---|---|---|---|
| | Mean | 95% CI | | Mean | 95% CI | | Mean | 95% CI | | |
| Acapulco, Guerrero | 3.633842 | 3.54619 | 3.71454 | 0.02521979 | 0.02175 | 0.02822 | 2.233462 | 2.16595 | 2.29749 | 22564 |
| Cabo San Lucas | 3.839943 | 3.73677 | 3.93137 | 0.03375925 | 0.02991 | 0.03702 | 2.110148 | 2.04734 | 2.17663 | 19798 |
| Monterrey, Nuevo Leon | 3.368576 | 3.25422 | 3.47348 | 0.0152396 | 0.01095 | 0.01932 | 2.177725 | 2.09471 | 2.26443 | 11992 |
| Juárez, Chihuahua | 3.316045 | 3.21866 | 3.39865 | 0.01497281 | 0.01147 | 0.01797 | 2.045822 | 1.97908 | 2.26443 | 17144 |
| Jojutla, Morelos | 3.701856 | 3.62331 | 3.77492 | 0.02699676 | 0.02416 | 0.02973 | 2.320845 | 2.24469 | 2.39917 | 17387 |
| Laboratory Population | 4.767385 | 4.55618 | 4.92226 | 0.0855144 | 0.07794 | 0.09134 | 1.627671 | 1.57480 | 1.68198 | 14068 |

**Table S5:** Data for adult survival on *Ae. aegypti* mosquitoes from Mexico along with one laboratory population were used to generate a Bayesian Weibull survival model. Mean and 95% credible interval (95% HPD interval) for the observed lifetimes ( $y_i$) at some particular temperature are drawn from a Weibull distribution with rate parameter( β ) and shape parameter( $k$). The Deviance Information Criterion (DIC) is given to compare model fits. We are comparing this model to one that fits all of the points together giving a DIC of 104,189.

**Supplementary Table 6:** Briere model output for biting rate for *Aedes aegypti* from Mexico compared to a laboratory adapted line.

| Population | $T_0$ | | | $T_m$ | | | *c* | | | DIC |
|---|---|---|---|---|---|---|---|---|---|---|
| | Mean | 95% CI | | Mean | 95% CI | | Mean | 95% CI | | |
| Acapulco, Guerrero | 10.21 | 6.94 | 12.47 | 36.99 | 36.12 | 38.25 | 1.93 e-04 | 1.50 e-04 | 2.31 e-04 | -693.32 |
| Cabo San Lucas | 9.61 | 5.13 | 12.68 | 39.65 | 37.36 | 43.42 | 1.76 e-04 | 1.21 e-04 | 2.31 e-04 | -559.26 |
| Monterrey, Nuevo Leon | 11.94 | 10.07 | 13.50 | 36.66 | 25.39 | 38.27 | 2.35 e-04 | 1.91 e-04 | 2.81 e-04 | -678.81 |
| Juárez, Chihuahua | 6.97 | 3.52 | 9.58 | 40.54 | 39.11 | 42.56 | 1.36 e-04 | 1.07 e-04 | 1.62 e-04 | -756.36 |
| Jojutla, Morelos | 5.32 | 0.72 | 9.88 | 41.19 | 38.28 | 44.43 | 1.27 e-04 | 9.31 e-05 | 1.76 e-04 | -608.79 |
| Laboratory Population | 7.7 | 0.87 | 13.62 | 40.19 | 35.98 | 44.64 | 1.55 e-04 | 9.47 e-05 | 2.53 e-04 | -338.90 |

| Population | Pmax | | | Topt | | | Breadth | | |
|---|---|---|---|---|---|---|---|---|---|
| | Mean | 95% CI | | Mean | 95% CI | | Mean | 95% CI | |
| Acapulco, Guerrero | 30.8 | 30.6 | 30.9 | 0.302 | 0.294 | 0.309 | 9.9 | 9.3 | 10.8 |
| Cabo San Lucas | 32.6 | 31.8 | 35.2 | 0.340 | 0.328 | 0.364 | 10.5 | 9.2 | 13.9 |
| Monterrey, Nuevo Leon | 30.7 | 30.3 | 31.6 | 0.326 | 0.319 | 0.336 | 9.3 | 8.4 | 10.8 |
| Juárez, Chihuahua | 33.1 | 32.6 | 34.3 | 0.316 | 0.308 | 0.327 | 11.7 | 10.8 | 13.7 |
| Jojutla, Morelos | 33.3 | 32.1 | 35.7 | 0.322 | 0.311 | 0.341 | 11.8 | 10.3 | 14.8 |
| Laboratory Population | 32.5 | 31 | 35.6 | 0.321 | 0.296 | 0.364 | 10.5 | 8.5 | 14.3 |

**Table S6:** Data for eggs per female per day taken from groups of mosquitoes across their lifetime for five populations of *Ae. aegypti* mosquitoes from Mexico compared to one laboratory adapted line were used to generate quadratic model. Mean and 95% credible interval (95% HPD interval) for the critical thermal minimum ($T_0$), maximum, ($T_m$), and a rate constant (*c*) are given for eggs per female per day. The Deviance Criterion Information (DIC) is given to compare model fits. We are comparing this model to one that fits all of the points together giving a DIC of -3382.774.

**Supplementary Table 7:** Quadratic model output for juvenile development rate for *Aedes aegypti* from Mexico compared to a laboratory adapted line.

| Population | a | | | b | | | *c* | | | DIC |
|---|---|---|---|---|---|---|---|---|---|---|
| | Mean | 95% CI | | Mean | 95% CI | | Mean | 95% CI | | |
| Acapulco, Guerrero | 1.72 e-03 | 1.06 e-03 | 2.13 e-03 | -7.44 e-02 | -9.31 e-02 | -4.16 e-02 | 0.769 | 0.368 | 0.987 | -73.48 |
| Cabo San Lucas | 4.48 e-04 | 1.16 e-04 | 9.21 e-04 | -1.91 e-02 | -4.41 e-02 | -2.70 e-03 | 2.15 e-01 | 1.70 e-02 | 5.22 e-01 | -72.815 |
| Monterrey, Nuevo Leon | 6.45 e-04 | 1.77 e-04 | 1.39 e-03 | -2.45 e-02 | -6.19 e-02 | -3.55 e-03 | 2.40 e-01 | 1.28 e-02 | 0.679 | -31.88 |
| Juárez, Chihuahua | 3.18 e-04 | 1.00 e-04 | 7.14 e-04 | -1.12 e-02 | -3.18 e-02 | -1.98 e-03 | 1.15 e-01 | 6.16 e-03 | 3.53 e-01 | -93.72 |
| Jojutla, Morelos | 6.12 e-04 | 1.13 e-04 | 1.30 e-03 | -2.56 e-02 | -6.03 e-02 | -2.93 e-03 | 2.66 e-01 | 1.40 e-02 | 0.689 | -32.93 |
| Laboratory Population | 5.93 e-04 | 6.95 e-05 | 1.42 e-03 | -2.69 e-02 | -6.85 e-02 | -1.92 e-03 | 3.28 e-01 | 2.81 e-02 | 0.819 | -33.02 |

**Table S7:** Data for juvenile mortality rate on *Ae. aegypti* mosquitoes from Mexico compared to a laboratory adapted line were used to generate quadratic model that is used in the fitness model. These data are from an approximation using larval survival rate and the amount of time to the adult stage. Mean and 95% credible interval (95% HPD interval) for the critical thermal minimum ($T_0$), maximum, ($T_m$), and a rate constant (*c*) are given for mosquito development rate. The Deviance Criterion Information (DIC) is given to compare model fits. We are comparing this model to one that fits all of the points together giving a DIC of -320.1.

**Supplementary Table 8:** Model output for temperature dependent fitness (*r*)

| Population | CTmin | | | CTmax | | | Topt | | |
|---|---|---|---|---|---|---|---|---|---|
| | Mean | 95% CI | | Mean | 95% CI | | Mean | 95% CI | |
| Acapulco, Guerrero | 19.68 | 18.3 | 21.2 | 33.4 | 32.3 | 34.1 | 26.1 | 26.1 | 27.6 |
| Cabo San Lucas | 18.41 | 17.3 | 19.3 | 33.04 | 32.4 | 33.7 | 28.21 | 27.8 | 28.6 |
| Monterrey, Nuevo Leon | 17.8 | 15.7 | 19.4 | 34.24 | 32.9 | 35.8 | 27.82 | 26.7 | 28.9 |
| Juárez, Chihuahua | 19.31 | 14.3 | 22.5 | 33.7 | 28.7 | 36.7 | 28.34 | 25.6 | 29.5 |
| Jojutla, Morelos | 18.78 | 13.9 | 22.2 | 31.11 | 26.0 | 35.0 | 26.84 | 23.7 | 28.6 |
| Laboratory Population | 18.36 | 17.2 | 19.3 | 33.67 | 32.6 | 34.7 | 27.65 | 26.8 | 28.4 |

| Population | Pmax | | | Breadth | | |
|---|---|---|---|---|---|---|
| | Mean | 95% CI | | Mean | 95% CI | |
| Acapulco, Guerrero | 0.2556054 | 0.2337234 | 0.2755966 | 6.2 | 6.0 | 6.2 |
| Cabo San Lucas | 0.5038521 | 0.4718735 | 0.5346916 | 7.9 | 7.4 | 8.3 |
| Monterrey, Nuevo Leon | 0.4560759 | 0.3785823 | 0.5329185 | 8.4 | 9.0 | 7.6 |
| Juárez, Chihuahua | 0.4048195 | 0.226706 | 0.4735402 | 7.5 | 3.7 | 8.9 |
| Jojutla, Morelos | 0.3583699 | 0.1547443 | 0.4844124 | 6.2 | 1.9 | 6.0 |

| | | | | | | |
|---|---|---|---|---|---|---|
| Laboratory Population | 0.4347708 | 0.3739203 | 0.4926068 | 7.9 | 7.2 | 8.4 |

**Table S8:** Data for mosquito temperature dependent population fitness on *Ae. aegypti* mosquitoes from Mexico along with one laboratory population were used to generate a composite model from mosquito development rate, eggs per female per gonotrophic cycle, adult survival, and juvenile mortality rate. Mean and 95% credible interval (95% HPD interval) for the critical thermal minimum (CTmin), maximum, (CTmax), optimum (Topt), thermal breadth (Tbreadth) and maximum thermal potential (Pmax) are given for fitness (*r*). We use the best fit model from each individual life history trait, making this the best fit model.

**Supplementary Table 9:** Temperature-dependent relative $R_0$ (transmission rate as the basic reproduction rate, $R_0$).

| Population | CTmin | | | CTmax | | | Topt | | |
|---|---|---|---|---|---|---|---|---|---|
| | Mean | 95% CI | | Mean | 95% CI | | Mean | 95% CI | |
| Acapulco, Guerrero | 20.2 | 23.1 | 18.7 | 32.77 | 31.3 | 33.7 | 28 | 27.9 | 28.1 |
| Cabo San Lucas | 17.8 | 23.0 | 17.7 | 33.8 | 32.3 | 33.6 | 28 | 28.2 | 27.9 |
| Monterrey, Nuevo Leon | 18.7 | 23.1 | 18.6 | 33.7 | 31.3 | 34.0 | 28.4 | 28.3 | 28.5 |
| Juárez, Chihuahua | 18.9 | 23.3 | 16.9 | 36.1 | 28.6 | 36.9 | 28.6 | 26.5 | 28.8 |
| Jojutla, Morelos | 17.4 | 16.6 | 23.0 | 34.2 | 35.1 | 26.0 | 26.93 | 24.7 | 27.8 |
| Laboratory Population | 18.1 | 23.0 | 17.1 | 34.7 | 32.5 | 34.9 | 26.1 | 27.1 | 25.9 |

| Population | Pmax | | | Breadth | | |
|---|---|---|---|---|---|---|
| | Mean | 95% CI | | Mean | 95% CI | |
| Acapulco, Guerrero | 1.0 | 0.7275 | 1.2772 | 4.9 | 4.1 | 5.2 |
| Cabo San Lucas | 1.0 | 0.7912065 | 1.232739 | 5.2 | 4.4 | 5.5 |
| Monterrey, Nuevo Leon | 1.0 | 0.7840912 | 1.243717 | 5.2 | 4.5 | 5.5 |
| Juárez, Chihuahua | 1.0 | 0.3312302 | 1.407316 | 5.1 | 2.6 | 5.7 |
| Jojutla, Morelos | 1.0 | 0.2240 | 1.6566 | 4.7 | 1.4 | 5.6 |
| Laboratory Population | 1.0 | 0.7361219 | 1.295784 | 5.4 | 4.3 | 6.1 |

**Table S9.** Temperature-dependent relative $R_0$ (transmission rate as the basic reproduction rate $R_0$) derived from individual life history traits measured for *Ae. aegypti* mosquitoes reared at 13°C, 15°C,19°C, 23°C, 25°C, 27°C, 29°C, 31°C, 33°C, 35°C, 37°C. We measured mosquito life history traits for five populations from Mexico (Cabo San Lucas, Acapulco, Monterrey, Ciudad Juárez, and Jojutla) compared to a laboratory-adapted line. Mean and 95% credible interval (95% HPD interval) for the critical thermal minimum (CTmin), maximum, (CTmax), optimum (Topt), thermal breadth (Tbreadth) and maximum thermal potential (Pmax) are given for fitness (*r*). We use the best fit model from each individual life history trait, making this the best fit model.
